# How a methanogen assimilates sulfate: Structural and functional elucidation of the complete sulfate-reduction pathway

**DOI:** 10.1101/2022.10.18.512691

**Authors:** Marion Jespersen, Tristan Wagner

## Abstract

By growing on sulfate as the sole source of sulfur, *Methanothermococcus thermolithotrophicus* breaks a dogma: the ancient metabolic pathways methanogenesis and sulfate-reduction should not co-occur in one organism due to toxic intermediates and energetic barriers. Using a complementary approach of physiological, biochemical, and structural studies, we provide a snapshot of the complete sulfate-reduction pathway of the methanogenic archaeon. While the first two reactions proceed via an ATP-sulfurylase and APS-kinase, common to other organisms, the further steps are catalysed by non-canonical enzymes. 3’-phosphoadenosine-5’-phosphosulfate (PAPS) released by the APS-kinase is converted into sulfite and 3’-phosphoadenosine-5’-phosphate (PAP) by a new class of PAPS-reductase that shares high similarity with the APS-reductases involved in dissimilatory sulfate-reduction. The generated PAP is efficiently hydrolysed by a PAP-phosphatase that was likely derived from an RNA exonuclease. Finally, the F_420_-dependent sulfite-reductase converts sulfite to sulfide for cellular assimilation. While metagenomic and metatranscriptomic studies suggest that genes of the sulfate-reduction pathway are present in various methanogens, *M. thermolithotrophicus* uses a distinct way to assimilate sulfate. We propose that its entire sulfate-assimilation pathway was derived from a “mix-and-match” strategy in which the methanogen acquired assimilatory and dissimilatory enzymes from other microorganisms and shaped them to fit its physiological needs.

## Introduction

Methanogens thrive in sulfidic habitats and receive sulfur through direct incorporation of environmental sulfides (HS^−^).^1,2^ In addition, there is accumulating evidence that some methanogens have the genomic potential to assimilate sulfur via sulfate reduction (SO_4_^2-^).^3^ To date, however, only one hydrogenotrophic methanogen has been experimentally shown to grow on SO_4_^2-^: *Methanothermococcus thermolithotrophicus*, a marine thermophile isolated from geothermally heated sea sediments close to Naples, Italy.^4,5^

Paradoxically, a methane-generating archaeon should not fix SO_4_^2-^ due to multiple obstacles. Firstly, methanogens commonly thrive in reduced environments where all other electron acceptors than CO_2_ are depleted, including SO_4_^2-^.^6,7^ Secondly, in the interface where methanogens and SO_4_^2-^ ions are co-occurring, hydrogenotrophic methanogens have to compete for the common substrate dihydrogen (H_2_) with dissimilatory SO_4_^2-^-reducing microorganisms. Thirdly, methanogens live at the thermodynamic limits of Life, and are considered energy extremophiles.^7,8^ Converting one SO_4_^2-^ molecule into HS^−^ requires 2-3 ATP, as well as reducing equivalents, a considerable investment for such energy-limited microorganisms.^4,7,8^ Lastly, the SO_4_^2-^ reduction pathway will generate the intermediate sulfite (SO_3_^2-^, or bisulfite HSO_3_^−^), which is a poison to the methane-forming reaction.^9^ But with the discovery of the F_420_-dependent sulfite-reductase (Fsr) it became evident that methanogens, such as *M. thermolithotrophicus*, which express Fsr can efficiently detoxify SO_3_^2-^ and even use it as sole sulfur source.^10-12^

The first step of the SO_4_^2-^ assimilation pathway is the capture and transport of the SO_4_^2-^-ion into the cell by an anionic transporter. Inside the cell, SO_4_^2-^ is activated by an ATP-sulfurylase (ATPS) to generate adenosine 5’-phosphosulfate (APS).^3,13,14^ From there, organisms can use different strategies (Supplementary Fig. 1, routes a-c): (1a) APS is directly reduced by an APS-reductase (APSR) to generate AMP and SO_3_^2-^. (1b) Alternatively, APS can be further phosphorylated to 3’-phosphoadenosine-5’-phosphosulfate (PAPS) by the APS-kinase (APSK). A PAPS-reductase (PAPSR) will reduce PAPS to SO_3_^2-^ and the toxic nucleotide 3’-phosphoadenosine 5’-phosphate (PAP). PAP must be quickly hydrolysed to AMP and inorganic phosphate by a PAP-phosphatase (PAPP). In both scenarios, the final step is carried out by a siroheme-containing sulfite-reductase, which reduces the SO_3_^2-^ into HS^−^. The latter can then be incorporated into biomass. (1c) In a different pathway, the sulfite-group of PAPS is transferred to another acceptor to build up sulfated metabolites. Route 1a is very similar to the dissimilatory pathway (Supplementary Fig. 1, route 2). However, dissimilatory APSRs and dissimilatory sulfite-reductases are structurally and phylogenetically distinct from their assimilatory counterparts and indirectly couple their reactions to membrane pumps, allowing for energy conservation.^15-17^

Genes encoding putative enzymes associated with SO_4_^2-^ reduction have been found in the genomes of multiple methanogens, including *M. thermolithotrophicus*.^3^ For this methanogen a theoretical, albeit incomplete, SO_4_^2-^ assimilation pathway can be hypothesized. Here, we elucidated the complete SO_4_^2-^ reduction machinery of this archaeon and describe how this one methanogen can convert SO_4_^2-^ into an elementary block of Life.

## Results

### SO_4_^2-^ growth dependency of *M. thermolithotrophicus*

Cultures grown on Na_2_S were successively transferred to a sulfur-free medium until no growth was observed. *M. thermolithotrophicus* showed robust growth when at least 100 μM of Na_2_SO_4_ were supplemented in the medium and reached similar cell yields as the Na_2_S grown culture. Under these cultivation conditions, SO_4_^2-^ is consumed over time as cell density increases (Fig. 1a). When cells are grown only on Na_2_S, no SO_4_^2-^ could be detected (Fig. 1a) indicating that *M. thermolithotrophicus* is not performing sulfide-oxidation.

**Fig. 1.**
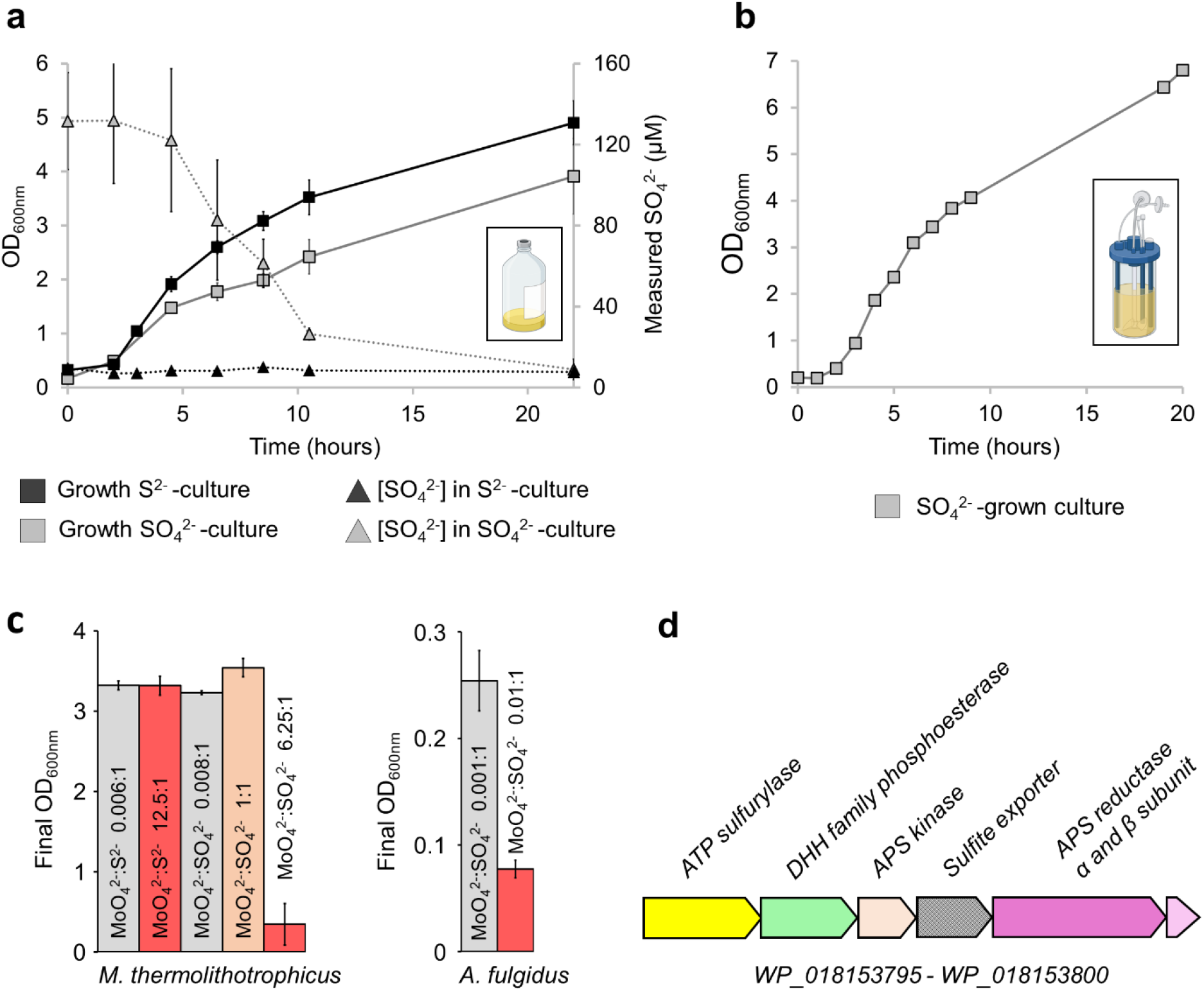
SO_4_^2-^-growth dependency of *Methanothermococcus thermolithotrophicus*. **a**, *M. thermolithotrophicus* cultures grown on Na_2_S (black squares, 0.5 mM) and Na_2_SO_4_ (grey squares, 0.5 mM). The consumption or release of SO_4_^2-^ in Na_2_S or Na_2_SO_4_ cultures are shown by black and grey triangles, respectively. All experiments were performed in triplicates and represented as data mean ± standard deviation (s.d.). Differences between expected (0.5 mM) and measured (0.13 mM) Na_2_SO_4_ concentration for the initial point are considered to be due to an artefact from the medium (see materials and methods). **b**, *M. thermolithotrophicus* grown on 10 mM Na_2_SO_4_ in a fermenter. The sampling points are visualized by grey squares. **c**, Molybdate (Na_2_MoO_4_) inhibition of Na_2_SO_4_ assimilatory and dissimilatory archaea. Growth experiments for *M. thermolithotrophicus* and *A. fulgidus* were performed in duplicates and quadruplicates, respectively. Results are represented as data mean ± s.d. **d**, Predicted operon for SO_4_^2-^ reduction from the whole genome shotgun sequence of *M. thermolithotrophicus*.

We then challenged the SO_42-_-grown culture by switching from batch to fermenter conditions, where H_2_S can escape to the gas phase and does not accumulate compared to flasks. In this open system with temperature and pH controlled, *M. thermolithotrophicus* grew to a maximum OD_600nm_ of 6.8 within 20 hours (Fig. 1b).

One way to decipher whether *M. thermolithotrophicus* relies on canonical enzymes of the SO_4_^2-^ reduction pathway is to use molybdate (MoO_4_^2-^). The structural analogue of SO_4_^2-^ binds to the ATPS and triggers molybdolysis, which hydrolyses ATP to AMP and pyrophosphate (PP_i_), resulting in cellular energy depletion.^18,19^ A MoO_4_^2-^:SO_4_^2-^ molar ratio of 0.004:1 is sufficient to inhibit the activity of dissimilatory SO_4_^2-^-reducing bacteria for 168 hours, an effect mainly due to molybdolysis by ATPS.^20-22^ SO_4_^2-^ assimilation is also affected by MoO_4_^2-^ as demonstrated by studies on plants.^23^ In the latter, growth inhibition occurred when MoO_4_^2-^ was in excess compared to SO_4_^2-^ and the ATPS activity was significantly affected at a 1:1 ratio.^19^ When applied on *M. thermolithotrophicus*, a high MoO_4_^2-^:Na_2_S ratio of 12.5:1 did not disturb growth of the Na_2_S-culture, indicating that MoO_4_^2-^ is not interfering with their basal metabolism. In contrast, a MoO_4_^2-^ :SO_4_^2-^ ratio of 6.25:1 was inhibitory to the SO_4_^2-^-grown culture, while a 1:1 ratio was not (Fig. 1c, Supplementary Fig. 2a). SO_4_^2-^ addition to the MoO_4_^2-^-inhibited culture restored growth (Supplementary Fig. 2b), indicating the reversibility of inhibition and its strict control by the MoO_4_^2-^:SO_4_^2-^ ratio rather than the MoO_4_^2-^ concentration. In comparison, in *Archaeoglobus fulgidus*, an archaeon that performs dissimilatory SO_4_^2-^ reduction to conserve energy, we observed growth inhibition at a MoO_4_^2-^:SO_4_^2-^ ratio of 0.01:1 (Fig. 1c, Supplementary Fig. 2c). These results suggest that *M. thermolithotrophicus* reduces SO_4_^2-^ via an assimilatory pathway containing a functional ATPS. Genes coding for putative standalone ATPS and APSK were indeed on the same locus in the genome of the strain DSM2095 that we had re-sequenced (Fig. 1d).^3,12,14^ To confirm their functions, the ATPS and APSK from *M. thermolithotrophicus* (*Mt*ATPS and *Mt*APSK, respectively) were further characterized.

### SO_4_^2-^ is activated and converted into PAPS by a conventional ATP-sulfurylase and APS-kinase

The activity of the recombinantly expressed *Mt*ATPS and *Mt*APSK was tested via a coupled assay (Fig. 2a, Supplementary Fig. 3) and a specific activity of 0.118 ± 0.008 μmol of oxidized NADH.min^−1^.mg^−1^ of *Mt*ATPS was measured. Under these conditions, the rate-limiting step was the pyrophosphatase activity. This highlights the need for rapid pyrophosphate degradation (Fig. 2a) to avoid a retro-inhibition as previously shown for other ATPS.^24^ A MoO_4_^2-^:SO_4_^2-^ ratio of 1:1.25 decreased the activity by half (see materials and method), corroborating that ATPS is also reacting with MoO_4_^2-^ as shown in other homologues.^19,22^

**Fig. 2.**
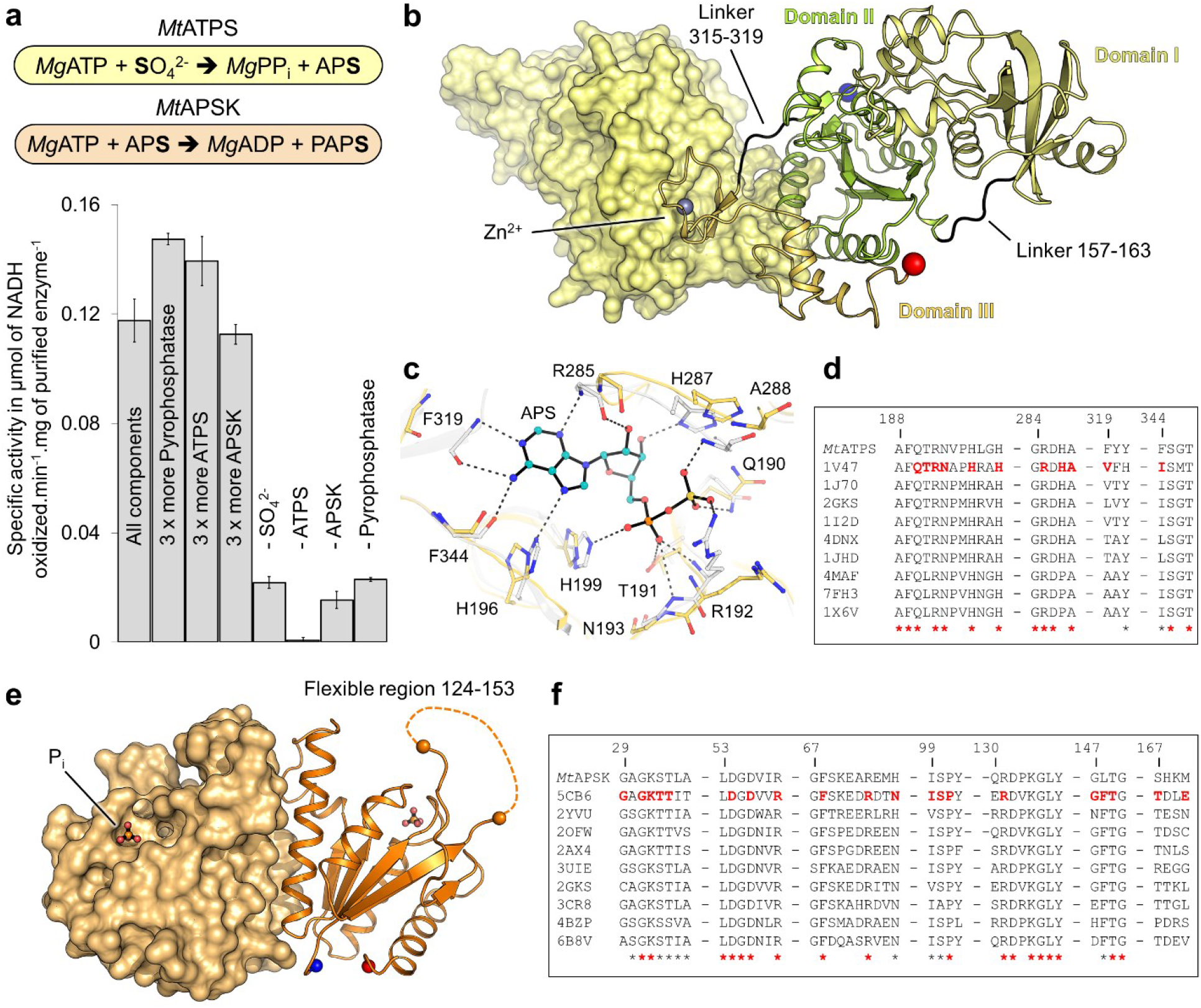
*Mt*ATPS and *Mt*APSK catalyse the first steps of the SO_4_^2-^ reduction pathway. **a**, The specific activity of *Mt*ATPS and *Mt*APSK, determined via a coupled enzyme assay. “-” indicates the absence of the respective reactant. All experiments were performed in triplicates and represented as data mean ± s.d. **b**, *Mt*ATPS homodimeric structure in which one monomer is shown as light yellow surface and the other one in cartoon. **c**, Active site of *Mt*ATPS superposed to the ATPS from *Thermus thermophilus* HB8 (PDB: 1V47, grey) containing the APS shown as balls and sticks. Residues involved in the substrate-binding are highlighted in sticks and only the ones from *Mt*ATPS are labelled. Nitrogen, oxygen, phosphorous and sulfur are coloured in blue, red, orange and yellow, respectively. **d**, Sequence conservation across ATPS homologues. **e**, *Mt*APSK homodimeric structure in which one monomer is shown as light orange surface and the other one in cartoon. The flexible loop, illustrated by dashed line could not be modelled. In all structures, the N- and C-terminus are shown by a blue and red sphere, respectively. **f**, Sequence conservation across APSK homologues. For d and f, red bold residues are involved in substrate binding while red and black stars are perfectly and well conserved residues, respectively.

The structure of *Mt*ATPS, the first representative of an archaeal ATPS, was refined to 1.97-Å resolution and obtained in an apo state despite co-crystallization with APS and SO_4_^2-^ (Supplementary Table 1). While the crystal packing suggests a homotetrameric assembly in two crystalline forms, size exclusion chromatography and surface analysis by the PISA server confirmed a homodimeric state, similar to bacterial homologues (Supplementary Fig. 4a and Supplementary Fig. 5). The structure exhibits the typical ATPS-fold comprised of three domains (domain I, 1-156; domain II, 164-314 and domain III, 320-382, Fig. 2b). The dimeric interface is mainly organized by the domain III as observed in *Thermus thermophilus*, a notable difference compared to other structural homologues (Supplementary Fig. 4a, b).^25-27^ Like in many thermophilic bacteria and archaea, the domain III contains a Zinc binding domain (320-343, Supplementary Fig. 4c, d) that might contribute to the thermal stability.^27^ *Mt*ATPS superposition with structural homologues shows a slight domain rearrangement probably due to the absence of substrate (Supplementary Fig. 4b). All residues critical for the reaction are conserved in *Mt*ATPS, arguing for a conserved reaction mechanism (Fig. 2c,d, Supplementary Fig. 4e, f and Supplementary Fig. 6).

The first APS-kinase model from a methanogen, *Mt*APSK, was refined to 1.77 Å. *Mt*APSK forms a homodimer with an organisation very similar to bacterial homologues (Supplementary Fig. 7a). Despite co-crystallization and soaking the crystals with APS and MgCl_2_, the *Mt*APSK structure was obtained in its apo state with a bound phosphate at the expected position of the ATP β-phosphate (Fig. 2e, Supplementary Fig. 7b, c). The N-terminus and region 124-153, the latter is involved in substrate binding^28,29^, could not be modelled due to the lack of electron density. However, the residues binding the substrates and Mg^2+^ are conserved (Fig. 2f, Supplementary Fig. 7b,c and Supplementary Fig. 8), suggesting that *Mt*APSK should be functional, as confirmed by the coupled enzyme assay.

### An alternative PAP-phosphatase with an exonuclease scaffold

If the ATPS and APSK are active they will produce PAPS, an intermediate which could follow the metabolic roads 1b or 1c (Supplementary Fig. 1). Both roads will lead to the production of the toxic product PAP, which inhibits sulfotransferases, exoribonucleases and disrupts the RNA catabolism.^30,31^ Therefore, it needs to be efficiently hydrolysed by a PAP-phosphatase. While the genome did not contain any related PAP-phosphatase, a gene coding for a putative phosphoesterase (Fig. 1d) was found in the genomic environment harbouring the ATPS and APSK genes. This PAP-phosphatase candidate, belonging to the DHH-family, was recombinantly expressed and produced inorganic phosphate (P_i_) from PAP at fast rates (5.1 μmol of P_i_ released.min^−1^.mg^−1^ of purified enzyme with manganese). The activity was stimulated by manganese addition and showed a high specificity towards PAP (Fig. 3a).

**Fig. 3.**
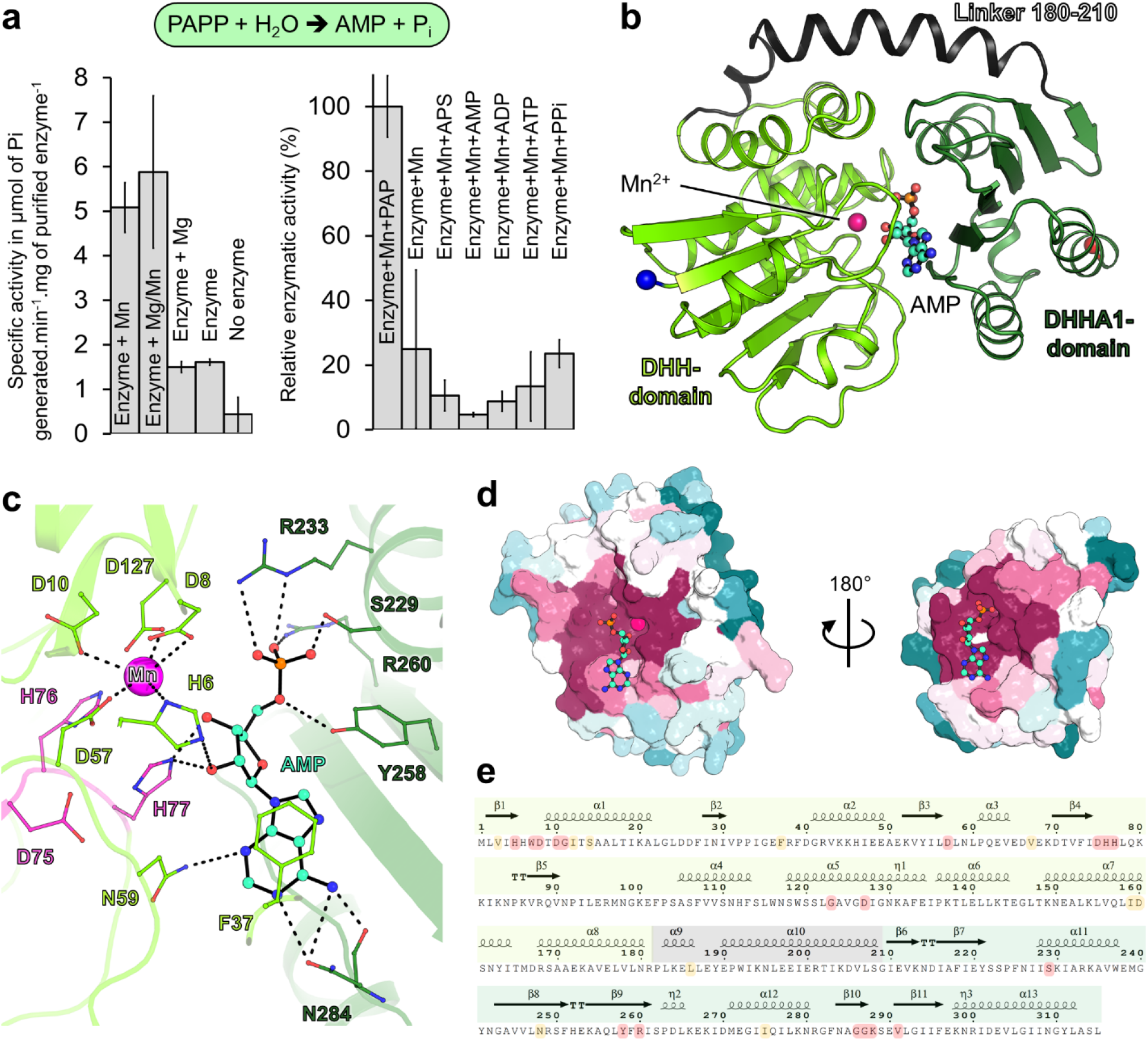
A new type of PAP-phosphatase. **a**, Specific activity of the *Mt*PAPP determined via the production of inorganic phosphate (P_i_) and relative enzymatic activity towards different nucleotides. All experiments were performed in triplicates and represented as data mean ± s.d. **b**, Organization of *Mt*PAPP shown in cartoon representation. The N- and C-terminus are highlighted as blue and red balls, respectively. Carbon, nitrogen, oxygen and phosphorous of AMP is coloured as cyan, blue, red and orange, respectively. **c**, Close-up of the active site of *Mt*PAPP. The residues coordinating the AMP and Mn^2+^ ion are highlighted by sticks and coloured as in b. **d**, Cut-through view of *Mt*PAPP structure shown in surface, and coloured by its sequence conservation across 168 archaeal homologues. The colour gradient ranges from variable (blue) to conserved (dark red). **e**, Secondary structure representation was done with ESPript 3.0^60^. The coloured frame corresponds to the different domains, DHH-domain in light green, linker in grey and DHHA1-domain in darker green. Perfectly conserved residues across 168 archaeal homologues are highlighted in red and the ones in yellow are well conserved.

To decipher the mechanism of this uncanonical PAP-phosphatase (baptised *Mt*PAPP), the enzyme was co-crystallized with manganese and PAP. The structure, solved by molecular replacement with a template generated by AlphaFold2^32,33^ was refined to 3.1-Å resolution and contained the product AMP and an ion in its active site, modelled as a partially occupied Mn^2+^ (Supplementary Table 1). *Mt*PAPP has no structural homologues but it shares an overall fold similar to the exonuclease RecJ or the oligoribonuclease NrnA from *B. subtilis* (*Bs*NrnA which also exhibits PAP-phosphatase activity, Supplementary Fig. 9a).^34-36^ The monomer is composed of a N-terminal (DHH, residues 1-179) and a C-terminal domain (DHHA1, residues 211-315) interconnected by a linker region (residues 180-210), forming a central groove (Fig. 3b). The DHH-domain contains the catalytic site and the DHHA1-domain serves as a scaffold to bind the substrate with high-specificity (Fig. 3b and c, Supplementary Fig. 9b). The motif coordinating the Mn^2+^ ion in RecJ and *Bs*NrnA is perfectly conserved in *Mt*PAPP^34,36^, therefore we expect that in its active state *Mt*PAPP would be loaded with two Mn^2+^. The first one, partially observed in the structure, is coordinated by four aspartates (Asp8, Asp10, Asp57, Asp127) and a long range interaction with His6. The absent second Mn^2+^ would be coordinated by the Asp10, Asp57, Asp127, the DHH motif (His76, His77) as well as by water molecules (Supplementary Fig. 9c). While the AMP shares a similar localization with structural homologues (β9β10β11), it is bound by a different interaction to the protein (Supplementary Fig. 9b, Supplementary Fig. 10). The nucleotide binding site would ideally place the 3′-phosphate of the PAP in front of the manganese when the enzyme is in its closed state (Supplementary Fig. 9c). The inter-domain movement, allowed by the linker, would facilitate a rapid exchange of the substrate/product, increasing the turnover of *Mt*PAPP. The complete sequence of this PAP-phosphatase was found in the genome of 168 archaea in which the nucleotide binding site is conserved (Fig. 3d-e, Supplementary Fig. 11). This suggests a common enzyme in archaea to detoxify PAP.

### An assimilatory PAPS-reductase structurally similar to dissimilatory APS-reductases

Which road uses *M. thermolithotrophicus* to metabolize PAPS to PAP: by a PAPSR-reductase (1b) or via a sulfo-transferase (1c)? No genes encoding these two enzymes were found in the genome, however, genes annotated as a dissimilatory APS-reductase (α- and β-subunit, APSR; Supplementary Fig. 12) are present and co-occur with the previously described genes (Fig. 1d). To experimentally confirm the activity and substrate specificity of this APS-reductase like enzyme, both subunits were co-expressed in *Escherichia coli*, purified and tested for enzyme activity assays (Fig. 4a). In contrast to dissimilatory APSRs, which catalyse the reversible reduction of APS to AMP and SO_3_^2-^,^37,38^ we could not measure the reverse reaction (i.e. AMP and SO_3_^2-^, or PAP and SO_3_^2-^ as substrates) for *M. thermolithotrophicus* enzyme by using K_3_Fe(CN)_6_ as an electron acceptor. Instead we used a coupled enzyme assay to reconstitute the pathway *in vitro* (Supplementary Fig. 13). *Mt*ATPS, a pyrophosphatase and the *Mt*APSK were used to generate PAPS. *Mt*PAPP was added to remove PAP, a potential retro-inhibitor of the reaction.^39^ The activity was monitored via the oxidation of reduced methyl viologen (MV_red_). When all components were present, a specific enzymatic activity of 0.114 ± 0.007 μmol of oxidized MV.min^−1^.mg^−1^ of the APS-reductase like enzyme was measured. A five time excess of the APS-reductase like enzyme resulted in a 220 % increase of the specific enzyme activity, indicating that the enzyme was the rate limiting step of the reaction (Supplementary Fig. 13). However, the accumulation of PAP (induced by the removal of *Mt*PAPP or Mn^2+^) strongly inhibited the activity. The specific enzymatic activity with APS as a substrate (i.e. removal of *Mt*APSK) was 0.007 ± 0.001 μmol of oxidized MV.min^−1^.mg^−1^ of the APS-reductase like enzyme (Fig. 4a). Considering the complexity of this coupled enzyme assay, kinetic parameters could not be determined but it did provide insights about the substrate specificity and confirmed that the APS-reductase like enzyme from *M. thermolithotrophicus* is exhibiting traits of a PAPS-reductase.

**Fig. 4.**
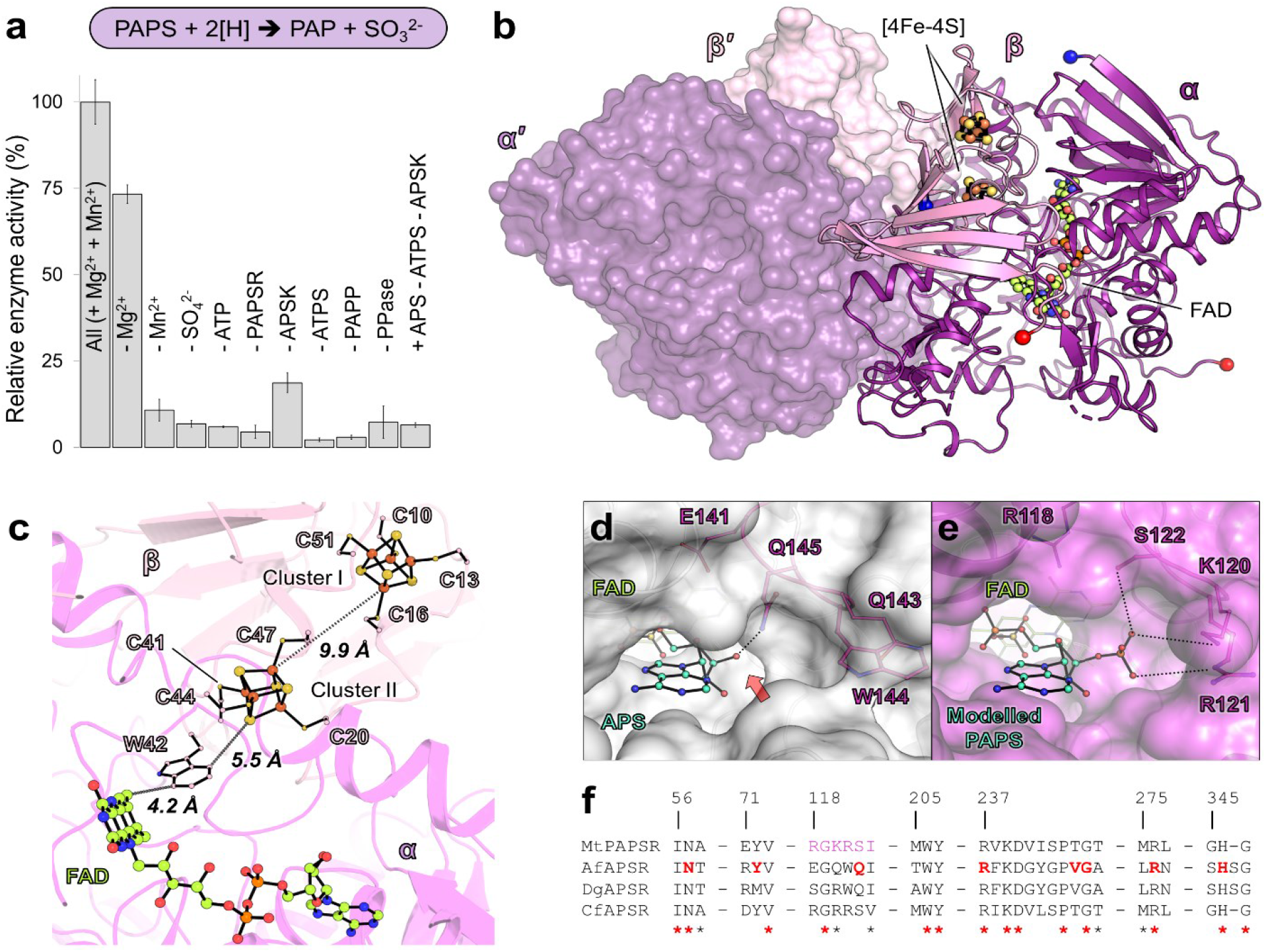
*Mt*PAPSR has a dissimilatory APS-reductase architecture but is specific to PAPS. **a**, Relative enzyme activity of *Mt*PAPSR, determined via a coupled enzyme assay (Supplementary Fig. 13). All experiments were performed in triplicates and represented as data mean ± s.d. **b**, *Mt*PAPSR organization with one heterodimer in surface and the other in cartoon. N- and C-terminus of both subunits are shown as balls and coloured in blue and red, respectively. Heterodimeric partners are labelled with a prime. Carbon, nitrogen, oxygen, phosphorous, iron and sulfur are coloured in limon, blue, red, orange, brown and yellow, respectively. **c**, Close-up of cofactors and the electron flow. [4Fe–4S]-clusters and cysteines coordinating them, FAD and the Trp42 proposed to participate in the electron transfer are shown in sticks and balls and coloured as in panel b. **d, e**, Active sites of APSR (d) from *A. fulgidus* containing APS (*Af*APSR, PDB: 2FJA) and *Mt*PAPSR with an artificially modelled PAPS (e) shown with a transparent surface. Residues involved in substrate recognition (based on modelled PAPS) are in balls and sticks and coloured as in panel b. A red arrow points to where PAPS would clash. **f**, Sequence conservation across the alpha subunit of *Mt*PAPSR, *Af*APSR, *D. gigas* (*Dg*APSR, PDB: 3GYX) and the putative APSR from *Caldanaerobius fijiensis* (*Cf*APSR, WP_073344903), which shares 68 % sequence identity with *Mt*PAPSR. Residues involved in APS binding for APSR are in bold and red, perfectly and well conserved residues are highlighted with a red and black stars, respectively. Trp206 and Tyr207 are involved in FAD binding. The sequence alignment was done with MUSCLE.^61^

To gain further molecular insights about the unconventional APS-reductase like from *M. thermolithotrophicus*, the enzyme was crystallized under anaerobic conditions. The structure was solved by a single-wavelength anomalous dispersion experiment measured at the Fe K-edge and refined to 1.45-Å resolution (Supplementary Table 1). The complex organizes as a α2β2 heterotetramer, with the same assembly than dissimilatory APS-reductases (Fig. 4b, Supplementary Fig. 14). It is, however, drastically different from characterized single-domain assimilatory APS/PAPS-reductases, which are thioredoxin/glutathione-dependent. They share no sequence or structural homology with *M. thermolithotrophicus* enzyme and several motifs that are proposed to mediate substrate binding and catalytic activity in assimilatory APS/PAPS-reductases are absent (Supplementary Fig. 14).^40^

In *M. thermolithotrophicus* the α-subunit, containing the FAD, is a member of the fumarate reductase family^37,41,42^ and the β-subunit is mainly composed of a ferredoxin-like domain in which two [4Fe–4S]-clusters are coordinated by eight cysteine residues (Fig. 4c). The residues coordinating APS, invariable in the dissimilatory family, differ in *M. thermolithotrophicus* and might provoke a switch of specificity from APS to PAPS. Despite a short soak with PAP, the putative substrate pocket contains only solvent, and we used the APS-reductase from *A. fulgidus* (*Af*APSR PDB: 2FJA, rmsd of 1.02-Å for 437 Cα aligned on the alpha subunit) to artificially model PAPS in the active site of the enzyme (Fig. 4e). The different substitutions mainly carried by the loop 104-123 would accommodate the additional 3′-phosphate-group by salt-bridge interactions and hydrogen bonds (Fig. 4d-f). In APS-reductases, however, a conserved glutamine (α145 in *A. fulgidus*) would clash with this phosphate group. The catalytic residues proposed in dissimilatory APS-reductases are retained in the enzyme of *M. thermolithotrophicus* (Supplementary Fig. 12, 14). We therefore propose an identical reaction mechanism based on a nucleophilic attack of the atom N5 of FAD on the sulfur PAPS, which creates a FAD-PAPS intermediate that decays to PAP and FAD-SO_3_^2-^.^37,42^ Taken together, the enzyme rates and the structural analysis, we propose that *M. thermolithotrophicus* harbours a new class of PAPS-reductase (*Mt*PAPSR) used to convert PAPS into SO_3_^2-^ and PAP.

### The last step of the pathway is catalysed by a F_420_H_2_-dependent sulfite-reductase

The SO_3_^2-^ generated by *Mt*PAPSR must be further reduced to HS^−^. In hydrogenotrophic methanogens, SO_3_^2-^ damages the methane-generating machinery and must be detoxified by the F_420_-dependent sulfite-reductase.^12^ We previously identified and characterized Group I Fsr in *M. thermolithotrophicus* (*Mt*Fsr) and determined a robust enzymatic activity towards SO_3_^2-^.^12^ Besides a second Fsr isoform, *M. thermolithotrophicus* does not contain other potential sulfite-reductases. While mass spectrometry confirmed that the Fsr isolated from SO_4_^2-^-grown cells is the characterized Group I *Mt*Fsr, the physiological role of the second Fsr isoform remains unknown.^12^ Therefore, *Mt*Fsr is the best candidate to catalyse the final reduction of SO_3_^2-^ to HS^−^. Native PAGE with cell extracts of cultures grown on different sulfur substrates confirmed the absence of *Mt*Fsr from cells grown on Na_2_S and its high abundance when grown on SO_3_^2-^.^10,12^

We determined a specific sulfite-reductase activity of 18.42 ± 0.13 μmol of oxidized MV.min^−1^.mg^−1^ of cell extract from Na_2_SO_3_ grown cells, in comparison to 7.31 ± 0.63 μmol of oxidized MV.min^−1^.mg^−1^ of cell extract from Na_2_SO_4_ grown cells, whereas cell extract of a Na_2_S grown culture had a specific sulfite-reductase activity of 3.04 ± 0.25 μmol of oxidized MV.min^−1^.mg^−1^ of cell extract (Fig. 5a). In agreement, we observed a band compatible with Fsr on the native PAGE for the SO_4_^2-^-grown culture but in lower amounts compared to SO_3_^2-^ conditions (Fig. 5b). The *Mt*Fsr structure recently published by our group was obtained from SO_4_^2-^ grown cells, which confirmed that it is the same enzyme expressed as under SO_3_^2-^ conditions.^12^ Taking together, these results argue that *Mt*Fsr is used as the last enzyme of the SO_4_^2-^ reduction pathway.

**Fig. 5.**
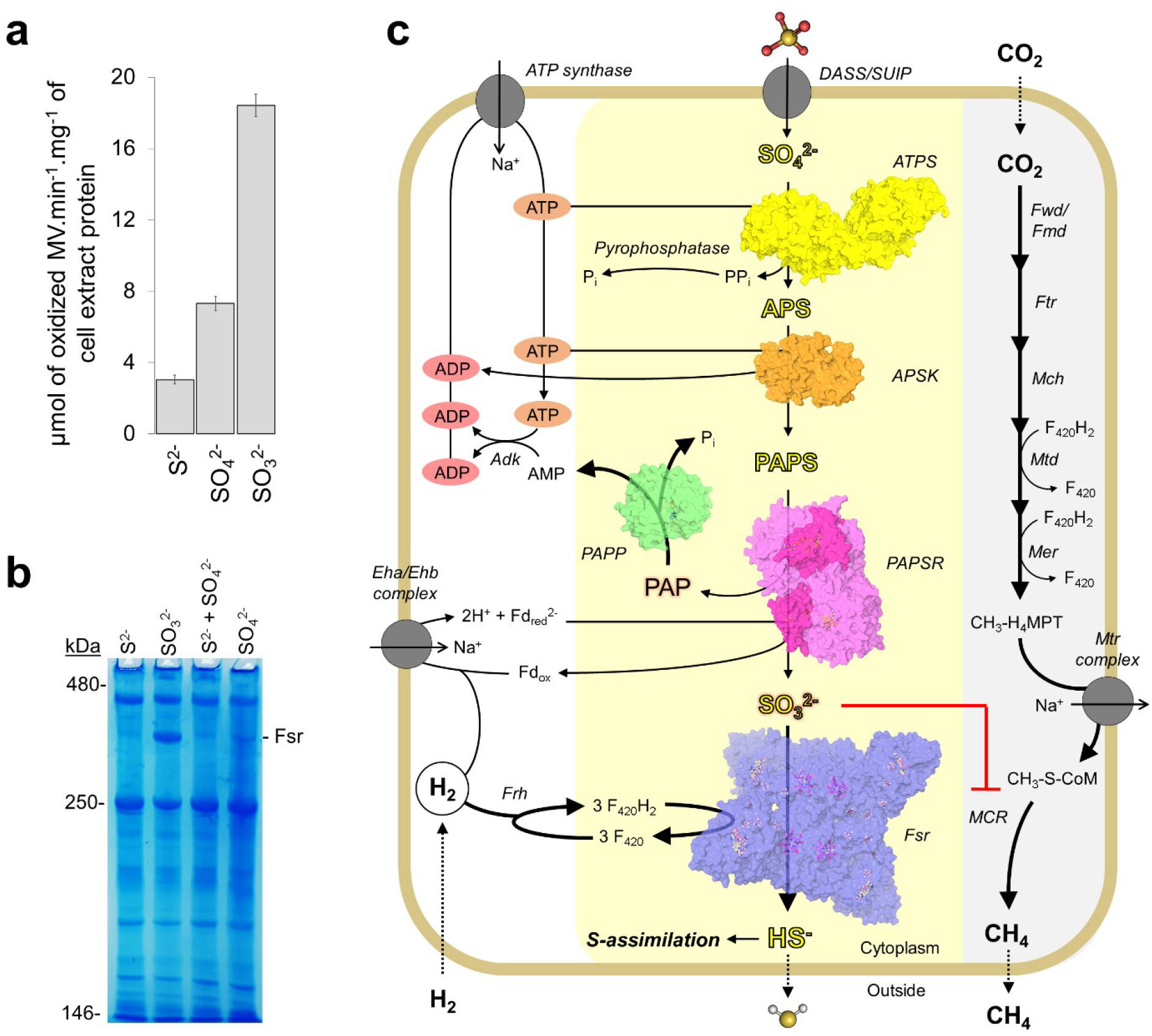
Proposed SO_4_^2-^ assimilation pathway in a methanogen. **a**, Sulfite-reductase activity in cell extract from *M. thermolithotrophicus* grown on different sulfur sources. All experiments were performed in triplicates and represented as data mean ± s.d. **b**, hrCN gel with *M. thermolithotrophicus* cell extract grown on different sulfur sources (15 μg protein loaded per sample). **c**, Proposed SO_4_^2-^ assimilation pathway in *M. thermolithotrophicus*. Yellow and grey backgrounds underline the SO_4_^2-^ reduction and methanogenesis pathways, respectively. Thick arrows indicate high metabolic fluxes. The structures of the enzymes operating the SO_4_^2-^ assimilation pathway are shown in surface with ligands as balls and sticks. Enzymes are abbreviated as follow: Fwd/Fmd, formylmethanofuran dehydrogenases; Ftr, tetrahydromethanopterin (H_4_MPT) formyltransferase; Mch, methenyl-H_4_MPT cyclohydrolase; Mtd, methylene-H_4_MPT dehydrogenase; Mer, 5,10-methylene-H_4_MPT reductase; Mtr, N^5^-CH_3_-H_4_MPT: coenzyme M methyl-transferase; Mcr, methyl-coenzyme M reductase; Adk, adenylate kinase; Frh, F_420_-reducing [NiFe]-hydrogenase; Eha/Ehb, energy-converting hydrogenase. Proposed candidates for DASS/SUIP putative SO_4_^2-^ transporters: WP_018154444/WP_018154062, respectively, pyrophosphatase: WP_018154062.

### Genetic potential is not enough to sustain growth on SO_4_^2-^: *Methanocaldococcus infernus* as a case study

Methanogens commonly use HS^−^ as a sulfur source, and the ones who express Fsr type I can also grow on SO_3_^2-^. Interestingly, some methanogens have genes which encode for proteins of the complete or partial SO_4_^2-^ reduction pathway.^3^ So why is *M. thermolithotrophicus* the only methanogen so far that has been proven to grow on SO_4_^2-^? We used *Methanocaldococcus infernus* as a model organism to investigate this further. *M. infernus* is a marine hyperthermophile that shares a very similar physiology than *M. thermolithotrophicus* and can grow in the same medium. It contains all genes coding for the enzymes characterized in this study with the exception of the described PAPSR. However, *M. infernus* genome encodes for a putative thioredoxin-dependent PAPSR and APSR, which share a high sequence identity with the biochemically characterized assimilatory APSR and PAPSR from *M. jannaschii* (Supplementary Fig. 15a).^43,44^ Therefore, *M. infernus* should, in principle, be capable of SO_4_^2-^ assimilation.

*M. thermolithotrophicus* and *M. infernus* were grown in the same medium and under the same cultivation conditions except that *M. infernus* was kept at 75 °C and *M. thermolithotrophicus* at 65 °C. *M. infernus* grew on 2 mM Na_2_S and Na_2_SO_3_ but was unable to use SO_4_^2-^ as a sole source of sulfur (Supplementary Fig. 15b).

This raises the question about the physiological function of the genes related to SO_4_^2-^ assimilation in methanogenic archaea. Based on our data, it could be that other methanogens still require these enzymes to acquire sulfur via the route 1c (Supplementary Fig. 1). The sulfur group would be transferred to an acceptor by a non-canonical sulfo-transferase, which might be important for uncharted biosynthetic pathway(s). That could explain why the PAPP gene is still present in methanogens containing the genes for ATPS and APSK. A counter-argument of this hypothesis is the presence of the thioredoxin-dependent PAPSR or APSR, characterized in *M. jannaschii*, which rather argues for route 1b.^3,43,44^ It is worth noticing that the gene coding for this putative assimilatory APSR also exists in *M. thermolithotrophicus* (WP_018154242.1). Therefore, further biochemical investigations will be needed to elucidate the physiological roles of those enzymes in methanogens.

## Discussion

This work unveiled the SO_4_^2-^ assimilation metabolism of one methanogenic archaeon, offering a molecular snapshot of the complete set of enzymes involved in the pathway. *M. thermolithotrophicus* activates SO_4_^2-^ by a conventional ATPS and APSK (Supplementary Fig. 16), but transforms it further by uncanonical enzymes (Fig. 5).

PAPS produced by the APSK is usually metabolized by thioredoxin- or glutathione-dependent assimilatory PAPS-reductases, which are organized as homo-oligomers. In contrast, *Mt*PAPSR inherited the heterotetrameric organization and FAD-based catalytic mechanism from dissimilatory APS-reductases (Fig. 4 and Supplementary Fig. 1, 14 and 17). We propose that the substitution of only a few amino acids switched the specificity towards PAPS (Fig. 4d-f), which might have been the result of a fine-tuned evolutionary adaptation to promote assimilatory SO_4_^2-^ reduction.

The generated PAP is efficiently hydrolysed by *Mt*PAPP. This PAP-phosphatase belongs to the DHH-family of phosphoesterases and shares structural homology with exonucleases. In comparison, conventional PAP-phosphatases (part of the FIG-superfamily), have a different fold (*i*.*e*. CysQ) and use three magnesium ions to hydrolyse the 3′-phosphate of PAP.^30^ *Mt*PAPP appears to be a remarkable example of convergent evolution that illustrates how archaea developed their own apparatus to efficiently detoxify PAP (Supplementary Fig. 17).

Group-I Fsr catalyses the final step of the SO_4_^2-^ reduction pathway. This enzyme shows distinct traits of dissimilatory sulfite-reductases with the active site composition of an assimilatory one.^12^ By encoding the *fsr* gene on a different locus, the methanogen is able to uncouple rapid SO_3_^2-^ detoxification from expressing the whole SO_4_^2-^ assimilation machinery. While the *Mt*ATPS, *Mt*APSK and *Mt*PAPSR show rather slow catalytic rates, *Mt*Fsr and the *Mt*PAPP have high specific activities, triggering the equilibrium towards HS^−^ production and efficiently eliminating toxic intermediates. Although our proposed pathway (Fig. 5c) would allow favourable thermodynamics, the first reactions should be regulated to avoid unnecessary ATP hydrolysis. We suspect that *Mt*ATPS, *Mt*APSK, and *Mt*PAPSR are cross-regulated by the accumulation of their own products, as already shown for homologues^39,45,46^, which would allow direct retro-control to harmonize the intracellular sulfur flux.

*M. thermolithotrophicus* is living at the thermodynamic limit of Life and the described SO_4_^2-^ assimilation will require the hydrolysis of three ATP to ADP for one processed SO_4_^2-^. While the methanogen could not bypass the ATP investment, it may have found an energy-saving strategy for the 8-electron reduction reaction from PAPS to HS^−^. Fsr uses F_420_, reduced by the F_420_-reducing hydrogenase, an adaptive strategy for a hydrogenotroph.^12,47^ Reduced F_420_ would be advantageous for *Mt*PAPSR, but it would require the assistance of a coupled F_420_H_2_-oxidase, which has not been identified yet. On the other hand, a reduced ferredoxin would be a suitable electron donor for this enzyme. Reduced ferredoxin could be obtained from the H_2_-dependent ferredoxin reduction via the Eha/Ehb complex, another advantageous strategy of hydrogenotrophs used for biosynthetic pathways (proposed in Fig. 5).

So far, it appears that the concomitant process of methanogenesis and complete SO_4_^2-^ reduction to HS^−^ is restricted to *M. thermolithotrophicus*, as it is the only methanogen that has been shown to grow on SO_4_^2-^. Strikingly, the only obvious difference between *M. thermolithotrophicus* and other methanogens that have the genomic potential to perform SO_4_^2-^ reduction is the acquisition of a PAPS-reductase belonging to the dissimilatory family (Supplementary Fig. 18). The physiological function of those SO_4_^2-^ reduction associated genes in other methanogens remains to be uncovered, as well as the benefit for *M. thermolithotrophicus* to assimilate SO_4_^2-^. From an ecological point of view, it might be beneficial, if not essential, for *M. thermolithotrophicus* survival to be capable of switching from H_2_S uptake to SO_4_^2-^ reduction. *M. thermolithotrophicus* was collected in geothermal heated sediments in a volcanic active area close to the shore of Naples. If the environment mimics a volcanic area, there could be soluble iron (Fe^2+^) that will react with the sulfide and with that deplete it as bioavailable sulfur. Volcanic sulfate aerosols or marine sulfate ions are then an alternative sulfur source. Such a fluctuating environment would explain the need to assimilate SO_4_^2-^.

The transplantation of *M. thermolithotrophicus* SO_4_^2-^ reduction system in methanogenic hosts which are already used as gas-converters (e.g. *Methanothermobacter*) would circumvent the requirements for highly toxic and explosive H_2_S through the inexpensive and abundant SO_4_^2-^. Beyond opening fantastic opportunities for safer biotechnological applications, a SO_4_^2-^-reducing hydrogenotrophic methanogen also reinforces the question whether sulfur-respiration or methanogenesis was the primeval means of energy-conservation during the evolution of early archaea. It has been previously suggested that methanogens are capable of dissimilatory-reduction of molecular sulfur at the expense of their methanogenesis metabolism.^48^ Both metabolisms are related to each other, and it is possible that the energetically more efficient methanogenesis originated from the sulfur-dissimilatory pathway. While most methanogens lost the ability to reduce SO_4_^2-^, some may have kept the required enzymes and adapted to an assimilatory pathway. However, this hypothesis is in conflict with the presented structural and phylogenetic analyses, since most enzymes of *M. thermolithotrophicus* SO_4_^2-^ assimilation machinery were likely acquired by horizontal gene transfer (Supplementary Fig. 16, 17)^49^. Therefore, it is more likely that *M. thermolithotrophicus* progressively assembled the entire SO_4_^2-^ reduction pathway via a “mix-and-match” strategy, providing a competitive advantage under fluctuating sulfur-source conditions and expanding its ecological niches.

## Materials & Methods

### Archaea strains and cultivation media

*M. thermolithotrophicus* (DSM 2095), *M. infernus* (DSM 11812) and *A. fulgidus* (DSM 4304) cells were obtained from the Leibniz Institute DSMZ-German Collection of Microorganisms and Cell Cultures (Braunschweig, Germany). *M. thermolithotrophicus* and *M. infernus* were cultivated in the same previously described minimal medium with some modifications.^12^ Refer to Supplementary information to find the complete composition of the media.

### Anaerobic growth of Archaea

The cell growth was followed spectrophotometrically by measuring the optical density at 600 nm (OD_600nm_). The purity of the culture was checked by light microscopy. The methanogens were cultivated with 1 × 10^5^ Pa of H_2_:CO_2_ with a 80:20 ratio in the gas phase. *M. infernus* was cultivated at 75 °C in 250 ml glass serum flasks and *M. thermolithotrophicus* was grown at 65 °C in flasks or fermenter. The serum flasks were not shaken but standing. *A. fulgidus* was cultivated in anaerobic and sealed 22 ml Hungate tubes, with 0.8 × 10^5^ Pa N_2_:CO_2_. 0.5 ml of the DSM 4304 culture was grown in 10 ml of classic media (see Supplementary information for media composition) containing 20 mM D/L-lactate. The culture was incubated at 80 °C, standing. All cultures were stored at room temperature in the dark under anaerobic conditions.

### Adaptation of *M. thermolithotrophicus* to SO_4_^2-^ and minimal SO_4_^2-^ requirement

*M. thermolithotrophicus* cells, grown on 2 mM Na_2_S, were successively transferred in 10 ml sulfur-free cultivation medium. After two transfers, the carry-over sulfur concentration of the inoculum did not support growth of *M. thermolithotrophicus*. By supplementing 2 mM Na_2_SO_4_ *M. thermolithotrophicus* growth resumed. No reducing agent was added to cope with the absence of HS^−^, which normally establishes a suitable reducing environment. Incubation without shaking is particularly important for reproducibility. Therefore, after inoculation, the cultures were incubated at 65 °C, standing for one night followed by shaking at 180 revolutions per minute (rpm) until they reached their maximum OD_600nm_. The gas phase was refreshed after the overnight incubation to maintain the pressure of 1 × 10^5^ Pa of H_2_:CO_2._ To measure the minimal SO_4_^2-^ concentration required to sustain growth, sulfur-limited *M. thermolithotrophicus* cells (using an inoculum to media ratio of 1:20) were provided with 2 mM, 1 mM, 0.5 mM, 0.25 mM, 0.1 mM and 0.04 mM Na_2_SO_4_. Growth was still observable for cells grown on 0.1 mM but not at 0.04 mM Na_2_SO_4_.

### SO_4_^2-^ measurements via Ion chromatography

Ion chromatography (Methrom Ion-Chromatograph) was used to measure the SO_4_^2-^ concentrations. A volume of 8 ml per sample was required with a maximum concentration of 0.5 mM SO_4_^2-^. SO_4_^2-^-reducing *M. thermolithotrophicus* cells were therefore grown in 1 L Duran bottles with 100 ml sulfur-free media, which was supplemented with 0.5 mM SO_4_^2-^ prior inoculation. As a negative control, 0.5 mM Na_2_S-grown *M. thermolithotrophicus* cells were used – inoculated and harvested similarly as the SO_4_^2-^-reducing cultures. All samples were taken aerobically and were passed through a 0.45 μM filter (Sartorius). If the cell densities were too high to be filtered, the samples were centrifuged at 13,000 x *g* for 7 minutes at 4 °C and the supernatant was taken for ion chromatography measurements. The samples were stored at 4 °C, when the measurements were not immediately performed.

### Growth of *M. thermolithotrophicus* in a fermenter

*M. thermolithotrophicus* was grown in a fermenter at 60 °C with 10 mM Na_2_SO_4_ as sole sulfur source. 7 L of anaerobic cultivation medium (see Sulfur-free cultivation medium for *Methanococcales*) supplemented with 10 mM Na_2_SO_4_ were continuously bubbled with H_2_:CO_2_ (80:20, 3 L.min^−1^). Under stirring (220 rpm), the medium was inoculated with 360 ml preculture (with an OD_600nm_ higher than 3). One hour after inoculation the culture was stirred at 800 rpm. 1 M NaOH was used as a base to readjust the pH upon acidification which was controlled using a pH probe. The cells were grown until late exponential phase (OD_600nm_ of 6.25 - 6.8) and then immediately transferred in an anaerobic tent (N_2_:CO_2_ atmosphere at a ratio of 90:10). Cells were harvested by anaerobic centrifugation for 30 min at 6,000 x *g* at 4 °C. Three SO_4_^2-^ -fermenter resulted in a final OD_600nm_ of 6.25, 6.42 and 6.8 after 20 hours of culture. 7 L of culture with an OD_600nm_ of 6.8 yielded 54 g of cells (wet weight). The cell pellet was transferred in a sealed bottle, gassed with 0.3 × 10^5^ Pa N_2_, flash frozen in liquid N_2_ and stored at -80 °C.

### Synthetic gene constructs

The DNA sequences of the ATP-sulfurylase, the APS-kinase, the PAP-phosphatase and the PAPS-reductase α and β subunits from *M. thermolithotrophicus* were codon optimized for *E. coli*, synthesized and cloned into pET-28a(+) vectors. For *Mt*ATPS, *Mt*APSK and the *Mt*PAPP the restriction sites NdeI and BamHI were used, with a stop codon (TGA) incorporated before BamHI. For *Mt*PAPSR a His-tag was placed at the C-terminus of the α-subunit, and a ribosome binding site was inserted between the coding sequences of the α- and β- subunits. The *Mt*PAPSR construct had the restriction sites NcoI and BamHI, with one stop codon incorporated after the His-tag for the α-subunit and one stop codon before BamHI for the β-subunit. These steps were performed by GenScript (GenScript Corp., Piscataway, NJ, USA). All sequences used are detailed in supplementary information.

### Enzyme overexpression and purification

All constructs were overexpressed and purified under aerobic conditions following a similar protocol, except for *Mt*PAPSR which was overexpressed and purified under an anaerobic atmosphere. All enzymes were passed on a HisTrap high performance column (GE healthcare, Sweden) followed, if necessary, by tag cleavage and gel filtration. See Supporting information for the complete protocol.

### Protein crystallization

Purified *Mt*ATPS, *Mt*APSK and *Mt*PAPP were kept in 25 mM Tris/HCl pH 7.6, 10 % v/v glycerol, 2 mM dithiothreitol and 150 mM NaCl. *Mt*PAPSR was kept in the same buffer without NaCl. Freshly prepared unfrozen samples were immediately used for crystallization. *Mt*ATPS, *Mt*APSK and *Mt*PAPP crystals were obtained under aerobic conditions at 18 °C. *Mt*PAPSR crystals were obtained anaerobically (N_2_:H_2_, gas ratio of 97:3), by initial screening at 20 °C. The sitting drop method was performed on 96-Well MRC 2-drop crystallization plates in polystyrene (SWISSCI) containing 90 μl of crystallization solution in the reservoir.

### Crystallization of *Mt*ATPS

*Mt*ATPS (0.7 μl) at a concentration of 14 mg.ml^−1^ (*Mt*ATPS form 1, Supplementary Table 1) or at a concentration of 27 mg.ml^−1^ (*Mt*ATPS form 2) was mixed with 0.7 μl reservoir solution. *Mt*ATPS at 27 mg.ml^−1^ was co-crystallized with 2 mM AMPcPP as well as 2 mM Na_2_SO_4_. For *Mt*ATPS form 1, transparent star-shaped crystals appeared after a few weeks in the following crystallization condition: 35 % w/v Pentaerythritol ethoxylate (15/4 EO/OH), and 100 mM 2-(*N*-morpholino)ethanesulfonic acid (MES) pH 6.5. For *Mt*ATPS form 2, transparent long but thin plate-shaped crystals appeared after a few weeks in the following crystallization condition: 20 % w/v polyethylene glycol 8000, 100 mM MES pH 6.0 and 200 mM Calcium acetate.

### Crystallization of *Mt*APS

*Mt*APSK (0.7 μl) at a concentration of 17.6 mg.ml^−1^ was mixed with 0.7 μl reservoir solution and co-crystallized with 2 mM MgCl_2_. Transparent, plate-shaped crystals appeared after a few weeks in the following crystallization condition: 20 % w/v polyethylene glycol 3350, and 100 mM tri-sodium citrate pH 5.5. *Mt*APSK was also crystallized with 2 mM MgCl_2_ and 2mM APS but the obtained structures of those crystals where of lower resolution and without any substrate or product present in the active site.

### Crystallization of *Mt*PAPSR

*Mt*PAPSR (0.7 μl) at a concentration of 20 mg.ml^−1^ was mixed with 0.7 μl reservoir solution and co-crystallized with FAD (0.5 mM final concentration). The crystal used for phasing was a brown flat-square and appeared after a few days in the following crystallization condition: 40 % v/v 2-Methyl-2,4-pentanediol, and 100 mM Tris/HCl, pH 8.0.

The crystal used to refine at high resolution was brown with an elongated plate-shape. It appeared after a few days in the following crystallization condition: 35 % v/v 2-Methyl-2,4-pentanediol, 100 mM Tris, pH 7.0 and 200 mM NaCl. Prior to transfer in liquid N_2_ the crystal was soaked in 10 mM disodium 3’-Phosphoadenosine 5’-phosphate for 7 min.

### Crystallization of *Mt*PAPP

*Mt*PAPP (0.7 μl) at a concentration of 20 mg.ml^−1^ was mixed with 0.7 μl reservoir solution and co-crystallized with Tb-Xo4 (10 mM final concentration), MnCl_2_ (2 mM final) and 2 mM PAP. The Tb-Xo4 is a nucleating/phasing agent^50^, which should increase the crystallization performance, however, in this case the same crystalline form has been obtained in absence of the compound and diffracted to similar resolution. Transparent, bipyramid crystals appeared after a few weeks in the following crystallization condition: 1.6 M tri-sodium citrate.

### X-ray crystallography and structural analysis

*Mt*PAPSR crystal handling was done inside the Coy tent under anaerobic atmosphere (N_2_:H_2_, 97:3), the other crystals were handled under aerobic conditions. The crystals were directly plunged in liquid nitrogen or were soaked for 5-30 seconds in their crystallization solution supplemented with a cryo-protectant prior to being frozen in liquid nitrogen. For *Mt*ATPS form 1, 30 % glycerol was used as cryoprotectant. For *Mt*APSK, 25 % ethylene glycol was used as cryoprotectant.

Crystals were tested and collected at 100 K at different synchrotrons (Supplementary Table 1). Data were processed with *autoPROC*^51^ except for *Mt*PAPP, which gave better statistics with indexation by XDS and scaling with *SCALA*^52^. All data collection statistics are mentioned in the Supplementary Table 1. *Mt*ATPS form 1 and 2, *Mt*APSK and *Mt*PAPP were solved by using *PHENIX* with the following templates: 1V47 (ATPS from *T. thermophilus*) for *Mt*ATPS form 1, *Mt*ATPS form 2 was solved using *Mt*ATPS form 1 as a template and *Mt*APSK was solved using 5CB6 (APS-kinase from *Synechocystis sp*.). For *Mt*PAPP the template was created de novo by AlphaFold 2.^32^

For *Mt*PAPSR, an X-ray fluorescence spectrum on the Fe-K edge was measured to optimize the data collection at the appropriate wavelength. Datasets were collected at 1.73646-Å for the single-wavelength anomalous dispersion experiment. Native datasets were collected at a wavelength of 0.97625-Å on another crystal. Data were processed and scaled with *autoPROC*^51^. Phasing, density modification and automatic building was performed with *CRANK-2*.^53^

All models were manually rebuilt with *COOT* and further refined with *PHENIX*.^54,55^ During the refinement, non-crystallographic symmetry and translational-liberation screw were applied. For all structures, except for ATPS form 1, hydrogens were added in riding position in the last refinement cycles. Hydrogens were removed in the final deposited models.

All models were validated through the MolProbity server.^56^ Data collection and refinement statistics, as well as PDB identification codes for the deposited models and structure factors are listed in the Supplementary Table 1. Figures were generated with PyMOL (Schrödinger, LLC). The metal in *Mt*ATPS was modelled as a Zinc based on CheckMyMetal server.^57^

### High resolution Clear Native PAGE (hrCN PAGE)

To visualize the expression levels of *Mt*Fsr when cells are grown with different sulfur sources, hrCN PAGE was performed. 10 ml of *M. thermolithotrophicus* cells supplemented with either 2 mM Na_2_S, 2 mM Na_2_SO_3_, 2 mM Na_2_S and 2 mM Na_2_SO_4_ as well as 2 mM Na_2_SO_4_ as sulfur substrates, were grown for one night at 65 °C, standing. Cells were harvested by anaerobic centrifugation at 6,000 x *g* for 20 min at room temperature and the cell pellets were resuspended in 2 ml lysis buffer (50 mM Tricine pH 8.0 and 2 mM sodium dithionite). The cells were sonicated 4 × 70 % intensity for 10 seconds, followed by a 30 second break (MS 73 probe, SONOPULS Bandelin, Germany). The hrCN PAGE was run anaerobically and the protocol was adapted from Lemaire et al.^58^

### Coupled enzyme activity of the *Mt*ATPS/*Mt*APSK

The activity of both enzymes was determined by the production of ADP which was coupled to NADH oxidation via the pyruvate kinase (PK) and the lactate dehydrogenase (LDH)^59^. The assays were performed in 96-deep well plates and spectrophotometrically monitored (Omega multi-mode Microplate Reader) at 360 nm at 35 °C. 100 mM KH_2_PO_4_ at pH 7.0, supplemented with 1.5 mM MgCl_2_ and 100 mM KCl, was used as a buffer. For NADH a molar extinction coefficient of 4,546.7 cm^−1^.M^−1^ was experimentally determined for the above-named conditions. To the buffer, 1 mM NADH, 2.5 mM Na_2_SO_4_, 1 mM PEP, 2 mM ATP, 2 U inorganic pyrophosphatase (*Saccharomyces cerevisiae*, 10108987001 from Sigma-Aldrich®), 1.1 U.ml^−1^ lactate dehydrogenase and 0.8 U.ml^−1^ pyruvate kinase (rabbit muscle, P0294 from Sigma-Aldrich®) and 0.5 mg.ml^−1^ *Mt*APSK (all final concentrations) were added. The reaction was started by the addition of 0.5 mg.ml^−1^ *Mt*ATPS. Addition of 0.02 mM Na_2_MoO_4_ did not affect activity (0.116 ± 0.027 μmol of oxidized NADH.min^−1^.mg^−1^) but the addition of 2 mM Na_2_MoO_4_ resulted in a decrease (0.068 ± 0.019 μmol of oxidized NADH.min^−1^.mg^−1^). All assays were performed in triplicates.

### *Mt*PAPP enzyme assay

The activity of the *Mt*PAPP was determined by the production of orthophosphate which was quantified using the Malachite Green Phosphate Assay Kit (Sigma-Aldrich) by forming a green complex. The assays were performed in 96-deep well plates and spectrophotometrically followed (Omega multi-mode Microplate Reader) by the absorbance at 620 nm. 25 mM Tris/HCl at pH 7.64 was used as a buffer. Buffer, 40 μM PAP or 90 μM of AMP/ADP/ATP/APS or PP_i_, 1 mg.ml^−1^ bovine serum albumin (BSA), 50 μM MnCl_2_ and / or 50 μM MgCl_2_ (final concentration) were mixed in a 1.5 ml Eppendorf tube one ice. 0.5 μg/ml of previously frozen *Mt*PAPP was added and the mixture was immediately incubated for 5 min at 40 °C. Next, 14 μl of the reaction mix was diluted in 66 μl of filtered Milli-Q H_2_O and immediately flash frozen in liquid N_2_ to quench the reaction. Then, 20 μl of malachite green reagent was added to the samples, the mixture was incubated at room temperature for 30 min and the formation of the green complex was measured at 620 nm. All assays were performed in triplicates. The measurements presented in Fig. 3 panel a come from two different experiments (left and right sub-panels). Both experiments were performed at two different days with the same enzyme preparation.

### Coupled *Mt*PAPSR assay

Since 3′-Phosphoadenosine-5′-phosphosulfate (PAPS) is unstable at high temperatures, we first tried to determine the activity of *Mt*PAPSR in the direction of PAPS production, as previously described for dissimilatory APS-reductases for APS production.^38^ PAPS oxidation was determined in 50 mM Tris/HCl buffer, pH 7.5, containing 5 mM Na_2_SO_3_, 2 mM PAP or 2 mM AMP (final concentrations) and 3.27 μg/ml *Mt*PAPSR. The reaction was started with a final concentration of 0.5 mM K_3_Fe(CN)_6_. The decrease in absorbance at 420 nm was measured and corrected for the background reaction without enzyme. No activity was detected.

Therefore we used the physiological reaction to monitor *Mt*PAPSR activity. In order to perform the coupled *Mt*PAPSR assay, the enzymes needed to be purified at the same time and immediately used for the assay. For each step, the *Mt*ATPS, *Mt*APSK and *Mt*PAPP were handled under aerobic conditions and on ice, while the *Mt*PAPSR was always kept in an anaerobic atmosphere and at 25 °C. To save time, the His-tags were not cleaved off. We previously saw that the tag did not interfere with the activity of these enzymes.

5.4 g, 7.6 g, 6.3 g and 11.0 g (wet weight) of recombinantly expressed *Mt*ATPS, *Mt*APSK, *Mt*PAPSR and *Mt*PAPP cells (as described above), respectively, were resuspended in 30 ml lysis buffer (50 mM Na_2_HPO_4_, pH 8.0, 500 mM NaCl, 20 mM Imidazole, 5 % Glycerol), separately. The cells were broken by the following sonication protocol: 5 cycles with 30 seconds at 75 % intensity followed by 1.5 min break (probe KE76, SONOPULS Bandelin) and cell debris were removed anaerobically via centrifugation (45,000 x *g*, 45 min at 4 °C). The filtered supernatant was applied on a Ni-NTA gravity column (1 ml Ni-NTA resin equilibrated with lysis buffer). The column was washed with 8 ml lysis buffer, then the proteins were eluted by applying 3 ml elution buffer (50 mM Na_2_HPO_4_, pH 8.0, 500 mM NaCl, 300 mM Imidazole, 5 % Glycerol). The flow trough was collected, filtered and injected onto a Superdex 200 Increase 10/300 GL (GE Healthcare), equilibrated in storage buffer (25 mM Tris/HCl pH 7.6, containing 10 % v/v glycerol and 2 mM DTT). The fractions containing the sample of interest were pooled and concentrated using 10 kDa (*Mt*PAPP) and 30 kDa cut-off centrifugal concentrator (Sartorius). The yield of this purification was: 2 ml of *Mt*ATPS at 11 mg.ml^−1^, 1.5 ml of *Mt*APSK at 8 mg.ml^−1^, 2 ml of the *Mt*PAPP at 3 mg.ml^−1^ and 2 ml of the *Mt*PAPSR at 4.5 mg.ml^−1^.

*Mt*PAPSR activity assays were carried out in an anaerobic atmosphere (100 % N_2_) at 45 °C. The assays were performed in 96-deep well plates and spectrophotometrically monitored on a SPECTROstar^®^ *Nano* Microplate Reader. 50 mM HEPES pH 7.0, supplemented with 50 mM KCl, 1.5 mM MnCl_2_ and 1.5 mM MgCl_2_, was used as a buffer. 0.5 mM reduced methyl viologen (MV_red_) served as an electron donor for *Mt*PAPSR. The molar extinction coefficient (ε_600nm_ = 8,133.3 cm^−1^·M^−1^) was experimentally determined using the above named conditions and by reducing methyl viologen with 2 mM sodium dithionite. For the assay, methyl viologen was reduced with carbon monoxide by the CO-dehydrogenase from *Clostridium autoethanogenum* according to the protocol from Lemaire and Wagner.^58^ CO was exchanged for N_2_ and the MV_red_ was immediately used for the assay. To the buffer and MV_red_, 5 mM ATP, 1 mM sodium dithionite, 0.2 U Pyrophosphatase (*E. coli*, MFCD00131379 from Sigma-Aldrich®), 0.127 mg.ml^−1^ *Mt*ATPS, 0.12 mg.ml^−1^ *Mt*APSK, 0.1 mg.ml^−1^ *Mt*PAPP and 0.064.5 mg.ml^−1^ *Mt*PAPSR were added. The reaction was started with the addition of 5 mM Na_2_SO_4_ and followed by oxidation of MV_red_ at 600 nm. All assays were performed in triplicates.

### Sulfite reductase activity in cell extracts

To determine the sulfite reductase activity from *M. thermolithotrophicus*, cultures were grown on either 2 mM Na_2_S, 2 mM Na_2_SO_3_ or Na_2_SO_4_ in 10 ml of the above mentioned medium in serum flasks. 9 ml of the cells were harvested in late exponential phase (OD_600nm_: 3.45 for 2 mM Na_2_S, 3.91 for 2 mM Na_2_SO_3_, 3.37 for Na_2_SO_4_) by centrifugation at 6,000 x *g* for 10 min at 4 °C. The supernatant was discarded and the cell pellets were frozen in liquid N_2_. The pellets were then resuspended in 1 ml 0.5 M KH_2_PO_4_ pH 7.0. The cells were lysed by sonication (2 × 10 seconds at 50 % intensity, probe MS73, SONOPULS Bandelin), followed by centrifugation at 4 °C at 15,600 x *g*. The supernatant was passed through a 0.2 μm filter and the protein concentration was determined by the Bradford method (6.63 mg.ml^−1^ for 2 mM Na_2_S, 6.14 mg.ml^−1^ for 2 mM Na_2_SO_3_, and 6.31 mg.ml^−1^ for Na_2_SO_4_). The activity assays were performed under an anaerobic atmosphere (100 % N_2_) at 50 °C, in 96-deep well plates and were spectrophotometrically monitored (SPECTROstar^®^ *Nano* Microplate Reader). The assay mixture contained 0.5 M KH_2_PO_4_ pH 7.0, 118 μM MV_red_ (final concentration, previously reduced with the equimolar amount of sodium dithionite) and 30 μM Na_2_SO_3_ (final concentration). Under those conditions a molar extinction coefficient of ε_600nm_ = 9,840 cm^−1^·M^−1^ was experimentally determined. The reaction was started by the addition of 0.05 μg of cell extract and followed by oxidation of MV_red_ at 600 nm. All assays were performed in triplicates.

### Phylogenetic trees

For a detailed description of the phylogenetic analysis refer to the supplementary data.

## Data Availability Statement

The structures were deposited in the protein data bank under the ID: 8A8G for *Mt*ATPS form 1, 8A8D for *Mt*ATPS form 2, 8A8H for *Mt*APSK, 8A8K for *Mt*PAPP and 8A8O for *Mt*PAPSR.

## Acknowledgements

We thank the Max Planck Institute for Marine Microbiology and the Max Planck Society for continuous support. We acknowledge the SOLEIL synchrotron for beam time allocation and the beamline staff of Proxima-1 for assistance with data collection. Furthermore, we thank the staff of beamline X06DA from SLS and P11 at PETRA III. We thank Prof. Dr Dennis R. Dean for providing us the RE plasmid pDB1281. We are thankful to Christina Probian and Ramona Appel for their continuous support in the Microbial Metabolism laboratory and cultivating *Archaeoglobus fulgidus*. We are grateful to Dr. Gunter Wegener and Martina Alisch from the HGF MPG Joint Research Group for Deep-Sea Ecology and Technology, for their assistance with the ion chromatography measurements. We deeply thank Dr. Ulrich Ermler, Prof. Dr. Julia Fritz-Steuber and Prof. Dr. Guenter Fritz for great discussions and critical comments regarding the manuscript.

## Author contributions

MJ cultivated the methanogens, purified and crystallized all proteins described in this study. MJ performed all biochemical characterization. MJ and TW collected X-ray data and solved the structures. MJ and TW refined all models and validated the models. TW and MJ designed the research and contributed to the writing of the article.

