## Supplementary information for "How a methanogen assimilates sulfate: Structural and functional elucidation of the complete sulfate-reduction pathway"

### Supplementary materials and methods.

Sulfur-free cultivation medium for *Methanococcales*. Per liter of medium: 558 mg of $KH_2PO_4$ (final concentration 4.1 mM), 1 g of KCl (13.4 mM), 25.13 g of NaCl (430 mM), 840 mg of $NaHCO_3$ (10 mM), 367.5 mg of $CaCl_2 \cdot 2\ H_2O$ (2.5 mM), 7.725 g of $MgCl_2 \cdot 6\ H_2O$ (38 mM), 1.18 g of $NH_4Cl$ (22.06 mM), 61.16 mg of nitrilotriacetic acid (0.32 mM), 6.16 mg of $FeCl_2 \cdot 4\ H_2O$ (0.031 mM), 10 µl of 2 mM $Na_2SeO_3 \cdot 5\ H_2O$ stock (0.02 µM), 3.3 mg of $Na_2WO_4 \cdot 2\ H_2O$ (0.01 mM) and 2.42 mg of $Na_2MoO_4 \cdot 2\ H_2O$ (0.01 mM) were dissolved under constant stirring with 750 ml of deionized $H_2O$ ($dH_2O$). 1 ml of 1.5 mM Resazurin and 10 ml of sulfur-free trace elements (see below) were subsequently added. In flasks the pH was set to either 7.6 with 50 mM Tris/HCl as buffer or to 6.2 with 50 mM 2-(*N*-morpholino)ethanesulfonic acid (MES). For the fermenter, 10 mM MES pH 6.2 was used as a buffer. The medium was filled up to a final volume of 1 L by the addition of $dH_2O$.

The cultivation media were transferred in a 1 L pressure protected DURAN® laboratory bottle with a magnetic stirring bar. The Duran flask was closed with a butyl rubber stopper and degassed by applying 3 min of vacuum, followed by 30 seconds addition of 1 X $10^5$ Pa $N_2$:$CO_2$ atmosphere (90:10), under constant magnetic stirring. This was repeated for a minimum of 15 cycles and at the final gas addition step, an overpressure of 0.3 X $10^5$ Pa $N_2$:$CO_2$ was applied.

Trace element composition for *Methanococcales*. A 100-fold-concentrated trace element solution was prepared by first dissolving 1.36 g of nitrilotriacetic acid (7.1 mM) in 800 ml $dH_2O$ under magnetic stirring. The pH was shifted to 6.2 by adding NaOH pellets. 89.06 mg of $MnCl_2 \cdot 4\ H_2O$ (0.45 mM), 183.3 mg of $FeCl_3 \cdot 6\ H_2O$ (0.68 mM), 60.27 mg of $CaCl_2 \cdot 2\ H_2O$ (0.41 mM), 180.8 mg of $CoCl_2 \cdot 6\ H_2O$ (0.76 mM), 90 mg of $ZnCl_2$ (0.66 mM), 37.64 mg of $CuCl_2$ (0.28 mM), 46 mg of $Na_2MoO_4 \cdot 2\ H_2O$ (0.19 mM), 90 mg of $NiCl_2 \cdot 6\ H_2O$ (0.38 mM) and 30 mg of $VCl_3$ (0.19

mM) was added separately. The trace element mixture was filled up to a final volume of 1 L with $dH_2O$.

Media for *Archaeoglobus fulgidus*. The media were modified from DSMZ media 399. It was prepared in a Widdel flask and contains, per liter of $dH_2O$: 0.14 g of $KH_2PO_4$, 0.25 g of $NH_4Cl$, 18 g of NaCl, 3.45 g of $MgSO_4 \cdot 7\ H_2O$, 4 g of $MgCl_2 \cdot 6\ H_2O$, 0.34 g of KCl, 0.14 g of $CaCl_2 \cdot 2\ H_2O$ and 1 ml of $Fe(NH_4)_2(SO_4)_2 \cdot 6\ H_2O$ (1.91 $mg.ml^{-1}$). The media was autoclaved for 25 minutes at 121 °C. Afterwards, the solution was transferred in a 1 L Duran bottle. 1 X $10^3$ Pa $N_2$:$CO_2$ (90:10) overpressure was applied during the addition of 1 ml of the trace elements M141 (see below), 0.05 mg of Vitamin $B_{12}$ (sterile filtered), 1 ml of a Se/Wo-solution (400 mg NaOH, 8 mg of $Na_2WO_4 \cdot 2\ H_2O$, 6 mg of $Na_2SeO_3 \cdot 5\ H_2O$ were solved in 1 L $dH_2O$ and autoclaved at 121 °C for 25 minutes), 1 ml of riboflavin (17.5 mM of acetic acid, 2.5 mg of riboflavin 5`-monophosphate sodium salt dihydrate were dissolved in 100 ml $dH_2O$, sterile filtered and stored in the dark at 4 °C), 0.1 ml of a thiamine solution (for a 100 ml solution 10 mg thiamine chloride hydrochloride were dissolved in 50 mM $Na_2HPO_4$/$H_3PO_4$ pH 3.7, sterile filtered and stored in autoclaved brown flasks at 4 °C until usage), 1 ml of 1 $mg.ml^{-1}$ Resazurin, 30 ml of 1 M $NaHCO_3$, and 1 ml of 5-vitamin mix (see below). For the $Na_2S$ grown cultures, 2 ml of 1 M $Na_2S$ and some crystals of sodium dithionite (until colour loss) were added but omitted for the $Na_2SO_4$-grown culture. The pH was adjusted to 6.9 using 2 M HCl.

Trace element composition for *Archaeoglobus fulgidus*. The 10 x trace element solution was modified from DSMZ media 141. 1.5 g nitrilotriacetic acid were dissolved in 80 ml Milli-Q® $H_2O$, and the pH was set to 6.5 using 1 M KOH. Subsequently 3 g of $MgSO_4 \cdot 7\ H_2O$, 0.5 g of $MnSO_4 \cdot H_2O$, 1 g of NaCl, 100 mg of $FeSO_4 \cdot 7\ H_2O$, 152 mg of $CoCl_2 \cdot 6\ H_2O$, 100 mg of $CaCl_2 \cdot 2\ H_2O$, 180 mg of $ZnSO_4 \cdot 7\ H_2O$, 10 mg of $CuSO_4 \cdot 5\ H_2O$, 20 mg of $KAl(SO_4)_2 \cdot 12\ H_2O$, 10 mg of $H_3BO_3$, 10 mg of $Na_2MoO_4 \cdot 2\ H_2O$ and 30 mg of $NiCl_2 \cdot 6\ H_2O$ were added and a pH 7.0 was set with 1 M KOH. The solution was filled up to 100 ml with Milli-Q® $H_2O$. The trace elements were autoclaved at 121 °C for 25 minutes and then stored at room temperature in the dark.

*Archaeoglobus fulgidus* 5 - Vitamin mix. 15 mg pyridoxine hydrochloride, 10 mg nicotinic acid, 5 mg calcium-D(+)-pantothenate, 4 mg 4-aminobenzoic acid, and 1 mg D(+)-biotin were dissolved in 100 ml 10 mM $Na_2HPO_4$ at pH 7.1, sterile filtered and stored at 4 °C until usage.

Protein overexpression. The *Mt*ATPS, *Mt*APSK and *Mt*PAPP constructs expressed in *Escherichia coli* strain BL21(DE3) were cultivated in 1 to 3 L of Lysogeny Broth (per liter of medium: 10 g tryptone, 5 g yeast extract, 10 g NaCl) supplemented with a final concentration of 50 µg/ml kanamycin. Cultures were incubated by shaking at 220 rotation per minute (rpm) at 37 °C until an $OD_{600nm}$ of 0.6 – 0.8 was reached. Induction was performed by adding a final concentration of 0.75 mM Isopropyl ß-D-1-thiogalactopyranoside (IPTG), and the cells were incubated for another hour by shaking at 37 °C. Cells were harvested by centrifugation at 5,000 × *g* for 20 min at 21 °C. Cell pellets were frozen in liquid $N_2$ and stored at -80 °C until further use.

For overexpression of the *Mt*PAPSR construct, *E. coli* BL21(DE3) was previously transformed with the plasmid pDB1282. The transformed cells were grown in a fermenter at 34 °C containing 8 L of modified Terrific Broth medium. For 8 L medium 96 g tryptone, 112 g yeast extract, 40 g glycerol, 4 g ferric ammonium citrate, 21 g MOPS pH 7.4 were dissolved in 7.2 L $dH_2O$ and autoclaved. Then 800 ml TB salts (10 x; 18.5 g $KH_2PO_4$, 100.32 g $K_2PO_4$, autoclaved) supplemented with glucose (28 mM final, sterile filtered), kanamycin (50 µg/ml final, sterile filtered), ampicillin (100 µg/ml final, sterile filtered) and riboflavin (20 µg/ml final), were added. The preculture was grown in classic Terrific Broth medium. The cells were gassed with a constant flow of 15 x $10^4$ Pa compressed air until they reached an $OD_{600nm}$ of 1.74, then the gas was switched to a constant flow of 15 x $10^4$ Pa $N_2$, to establish anaerobic conditions. Next, the cells were induced with a final concentration of 0.2 % L-arabinose, 25 mM sodium fumarate dibasic, 2 mM cysteine hydrochloride and 50 µM IPTG followed by incubation for one hour at 28 °C. The cells were harvested under anaerobic conditions by centrifugation at 5,000 × *g* for 20 min at 4 °C. Cell pellets were frozen in liquid $N_2$ and stored at -80 °C until further use.

*Mt*ATPS, *Mt*APSK and *Mt*PAPP tag-cleavage and purification. 6 g, 9 g and 7 g (wet weight) of respectively *Mt*ATPS, *Mt*APSK, *Mt*PAPP-overexpressed *E.coli* cells were thawed under warm water and were resuspended in 22-50 ml lysis buffer (50 mM $Na_2HPO_4$ pH 8.0, 500 mM NaCl, 20 mM Imidazole, 5 % Glycerol) on ice. The cell lysate was homogenized by sonication: 10 cycles with 1 min at 80 % intensity followed by 1.5 min break (probe KE76, SONOPULS Bandelin) and cell debris were removed via centrifugation (45,000 x *g*, 50 min at 4 °C). The filtered sample was applied to a 5 ml HisTrap high performance column (GE healthcare, Germany), which was previously equilibrated with lysis buffer. The column was then washed with 2 column volumes of lysis buffer. A gradient of 0.02 to 0.3 M Imidazole was applied for 40 min at a flow rate of 1.5 $ml.min^{-1}$ and fractions of 1 ml were collected. *Mt*ATPS eluted between 0.1 and 0.15 M Imidazole, *Mt*APSK between 0.12 and 0.2 M and *Mt*PAPP eluted between 0.13 and 0.22 M Imidazole. The protein fractions were pooled and the buffer was exchanged for Phosphate-buffered saline (137 mM NaCl, 2.7 mM KCl, 10 mM $Na_2HPO_4$, 1.8 mM $KH_2PO_4$ pH 7.4) by using 30 kDa cutoff filter (6 ml, Merck Millipore, Darmstadt, Germany) for *Mt*ATPS and *Mt*APSK and a 10 kDa cutoff filter for *Mt*PAPP. The proteins were concentrated to 3 ml for the *Mt*PAPP and 5 ml for the *Mt*ATPS and *Mt*APSK. The *Mt*PAPP with the His-Tag was immediately passed onto a Superdex 200 Increase 10/300 GL (GE Healthcare), equilibrated in storage buffer (25 mM Tris/HCl pH 7.6, containing 5 % v/v glycerol and 2 mM dithiothreitol), as the protein aggregated with time. *Mt*PAPP eluted at a flow rate of 0.8 $ml.min^{-1}$ in a sharp Gaussian peak at an elution volume of 75 ml. The fractions of interest containing *Mt*PAPP were concentrated with a 10 kDa cutoff centrifugal concentrator to 150 µl and the protein was directly used for crystallization. The concentration of purified *Mt*PAPP, estimated by the Bradford method, was 20 $mg.ml^{-1}$.

For the *Mt*ATPS and *Mt*APSK, 50 µl of 0.1 $U.mg^{-1}$ Thrombin (from bovine plasma, Sigma-Aldrich, Germany) was added to the 5 ml of protein and incubated overnight at 22 °C to remove the His-Tag of the enzymes. The samples were then passed onto a HiLoad® 16/600 Superdex® 200 pg (GE Healthcare) equilibrated in storage buffer (25 mM Tris/HCl pH 7.6, containing 150 mM NaCl, 10 % v/v glycerol and 2 mM dithiothreitol). *Mt*ATPS and *Mt*APSK eluted at a flow rate of 0.8 $ml.min^{-1}$ in a sharp Gaussian peak at an elution volume of 68 ml and 81 ml, respectively. The fractions of interest containing the proteins were concentrated with a 30 kDa cutoff centrifugal concentrator (6 ml, Merck Millipore, Darmstadt, Germany) to 150 µl and the proteins were immediately used for crystallization. The concentration of purified *Mt*ATPS, estimated by the Bradford method, was 27 $mg.ml^{-1}$ and 17.6 $mg.ml^{-1}$ for *Mt*APSK.

<u>*Mt*PAPS-reductase purification.</u> 26 g (wet weight) of *Mt*PAPSR-overexpressed *E.coli* cells from the fermenter were thawed under warm water and transferred to an anaerobic tent containing an atmosphere of $N_2$:$CO_2$ (with a 90:10 ratio). 120 ml lysis buffer (50 mM $Na_2HPO_4$, 500 mM NaCl, 20 mM Imidazole, 2 mM dithiothreitol, 5 % Glycerol) was added and cells were lysed by sonication: 5 cycles with 1 min at 75 % intensity followed by 3 min break (probe KE76, SONOPULS Bandelin). Cell debris were removed anaerobically via centrifugation (45,000 x *g*, 45 min at 4 °C). The supernatant was transferred to a Coy tent ($N_2$:$H_2$ atmosphere with a 97:3 ratio) under yellow light at 20 °C and filtered through a 0.2 µm filter (Sartorius). The filtered sample was applied to a 5 ml HisTrap high performance column (GE healthcare), which was previously equilibrated with lysis buffer. The column was then washed with 2 column volumes of lysis buffer. A gradient of 0.02 to 0.3 M Imidazole was applied for 40 min at a flow rate of 1.5 $ml.min^{-1}$ and fractions of 1 ml were collected. *Mt*PAPSR eluted between 0.04 and 0.15 M Imidazole. The fractions of interest were merged and diluted with 4 volumes of 50 mM Tricine/NaOH pH 8.0 and 2 mM dithiothreitol. The sample was filtered through 0.2 µm and was loaded on a 5 ml Q Sepharose high performance column (GE healthcare). A gradient of 0 to 0.55 M NaCl was applied for 90 min with a flow rate of 1 $ml.min^{-1}$. Fractions of 1.5 ml were collected. *Mt*PAPSR eluted between 0.11 and 0.52 M NaCl.

The purest *Mt*PAPSR fractions were pooled and the buffer was exchanged for storage buffer (25 mM Tris/HCl pH 7.6, containing 10 % v/v glycerol and 2 mM dithiothreitol) by using 30 kDa cutoff filter (6 ml, Merck Millipore, Darmstadt, Germany) and *Mt*PAPSR was concentrated to 400 µl. The concentrated sample was passed onto a Superdex 200 Increase 10/300 GL (GE Healthcare), equilibrated in storage buffer. *Mt*PAPSR eluted at a flow rate 0.4 $ml.min^{-1}$ in a sharp Gaussian peak at an elution volume of 12.5 ml. The fractions of interest containing *Mt*PAPSR were concentrated with a 30 kDa cutoff centrifugal concentrator to 300 µl and the protein was directly used for crystallization. For the activity assays *Mt*PAPSR was incubated with 0.5 mM FAD for 15 min to promote cofactor integrity, followed by buffer exchange to remove excess FAD using a 30 kDa cutoff concentrator. The concentration of purified *Mt*PAPSR, estimated by the Bradford method, was 20 $mg.ml^{-1}$.

<u>Phylogenetic trees</u>. Phylogenetic analyses were performed using MEGA11 by applying default parameters.[1] Homolog proteins were first identified and obtained from NCBI using BLASTP (E-value cut-off of 1e1). The protein sequences were then aligned using MUSCLE and all homologs were identified through an iterative alignment evaluation based on characterized proteins and manual filtering. The evolutionary history was inferred using the Neighbor-Joining method. The

bootstrap consensus tree inferred from 2000 replicates is taken to represent the evolutionary history of the taxa analysed. Branches corresponding to partitions reproduced in less than 50 % bootstrap replicates are collapsed. The percentage of replicate trees in which the associated taxa clustered together in the bootstrap test (2000 replicates) are shown next to the branches. The evolutionary distances were computed using the JTT matrix-based method and are in the units of the number of amino acid substitutions per site.

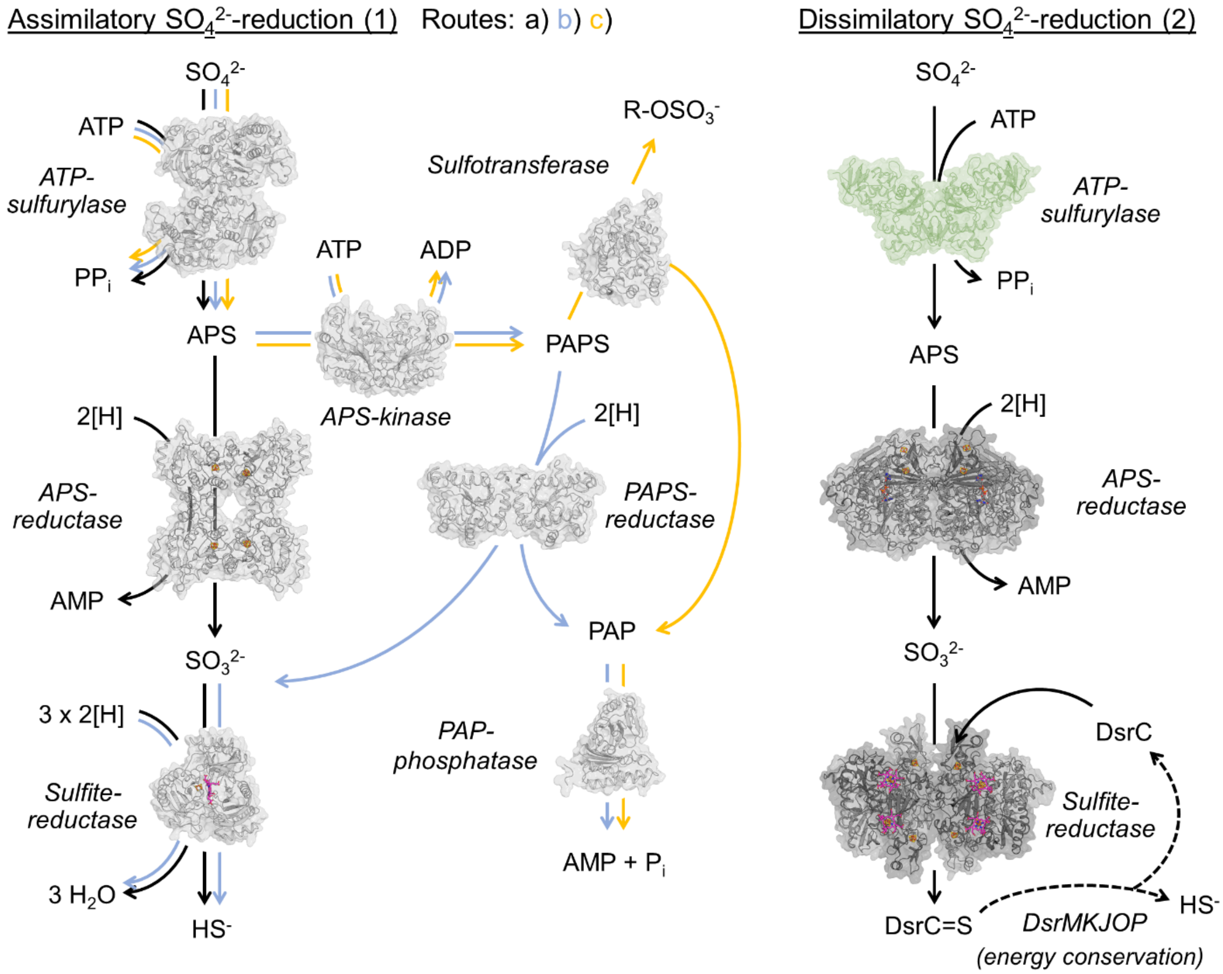


**Supplementary Fig 1. Sulfate ($SO_4^{2-}$)-reduction pathways.** Assimilatory $SO_4^{2-}$-reduction. (Route 1a, 1b, 1c) $SO_4^{2-}$ is activated by the ATP-sulfurylase (e.g. *Glycine max*, PDB: 4MAF) to APS. (1a) APS gets directly reduced to sulfite ($SO_3^{2-}$) and AMP by a one [4Fe-4S]-cluster containing APS-reductase (e.g. *Pseudomonas aeruginosa*, PDB: 2GOY, thioredoxin dependent). (1b, c) Alternatively, the APS gets further phosphorylated by an APS-kinase (e.g. *Arabidopsis thaliana*, PDB: 3UIE) to produce PAPS. (1b) A PAPS-reductase (*Saccharomyces cerevisiae*, PDB: 2OQ2, thioredoxin dependent) converts PAPS into $SO_3^{2-}$ and PAP. The PAP will be hydrolysed to inorganic phosphate ($P_i$) and AMP by a PAP-phosphatase (e.g. *Mycobacterium tuberculosis*, PDB: 5DJJ). (1a, b) A sulfite-reductase (*Escherichia coli*, PDB: 1AOP) reduces the $SO_3^{2-}$ into $S^{2-}$, which can then be incorporated into biomass. In the route 1c, a sulfotransferase (e.g. *A. thaliana*, PDB: 5MEK) catalyses the transfer of the sulfo-group (R-$OSO_3^-$) from PAPS to an alcohol or amine acceptor. Dissimilatory $SO_4^{2-}$-reduction. $SO_4^{2-}$ is activated by the ATPS to APS and further reduced to $SO_3^{2-}$ by an APS-reductase (e.g. *Archaeoglobus fulgidus*, PDB: 2FJA), which contains two [4Fe-4S]-cluster. A sulfite-reductase (e.g. *A. fulgidus*, PDB: 3MM5) reduces the $SO_3^{2-}$ and branches it on the carrier DsrC. The membrane complex DsrMKJOP (e.g. *Allochromatium vinosum*) reduces the sulfur into $HS^-$ concomitantly with ion translocation for energy conservation. Sirohemes and [4Fe-4S]-clusters are represented in sticks and spheres, with carbon, oxygen, nitrogen, sulfur and iron coloured in pink, red, blue, yellow and orange. The enzymes are shown in cartoon and transparent surface in their oligomeric state. The ATPS of *A. fulgidus* was modelled using Alphafold2[2] and coloured in green. The bifunctional ATP-sulfurylase CysDN using an additional GTP, was not presented here to simplify the scheme.

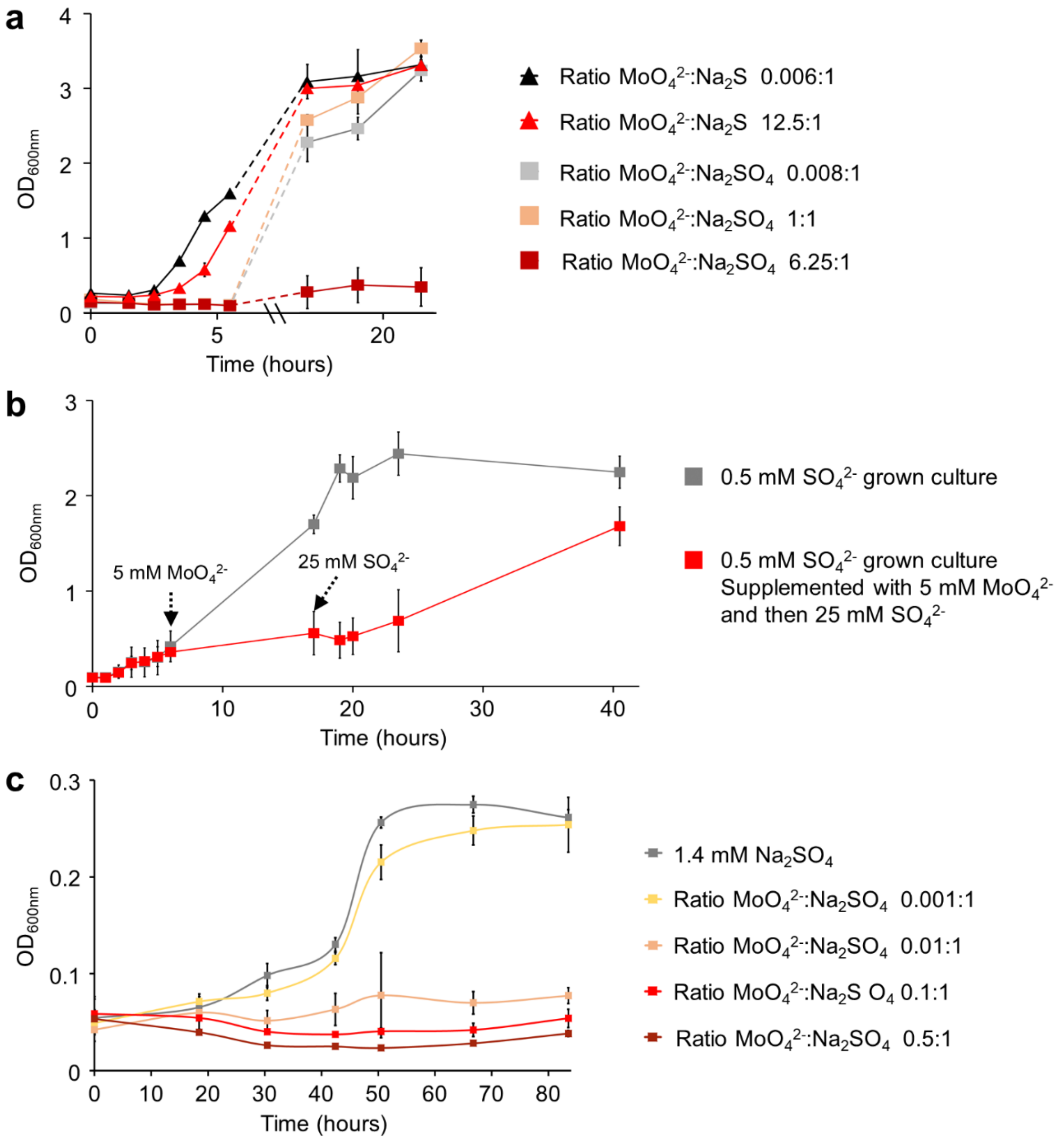


**Supplementary Fig 2. Impact of Molybdate ($MoO_4^{2-}$) on $SO_4^{2-}$-reducing archaea. a**, $MoO_4^{2-}$ tolerance of *M. thermolithotrophicus*. The archaeon was grown on $SO_4^{2-}$ (grey square), $SO_4^{2-}$ supplemented with an equimolar amount of $MoO_4^{2-}$ (wheat square) and $SO_4^{2-}$ supplemented with an excess of $MoO_4^{2-}$ (dark red square). As a control, $Na_2S$ grown cultures ($S^{2-}$, black triangle) and $Na_2S$ grown cultures with an excess of $MoO_4^{2-}$ (red triangle) were used. This growth experiment was performed in duplicates. **b**, Effect of $MoO_4^{2-}$ : $SO_4^{2-}$ ratios in *M. thermolithotrophicus* cultures grown on 0.5 mM $Na_2SO_4$. Grey squares indicate the growth curve of $SO_4^{2-}$-reducers without addition of $MoO_4^{2-}$. The red squares indicate the growth curve of $SO_4^{2-}$-reducers exposed to 5 mM of $MoO_4^{2-}$, followed by the addition of 25 mM $Na_2SO_4$. Black, dashed arrows indicate time of $MoO_4^{2-}$ and $SO_4^{2-}$ addition. This growth experiment was performed in triplicates. **c**, *Archaeoglobus fulgidus* sensitivity towards $MoO_4^{2-}$. Here, a $MoO_4^{2-}$:$SO_4^{2-}$ ratio of 0.01:1 is sufficient to inhibit growth of *A. fulgidus*. The data shown are quadruplicates except for the lowest and the highest $MoO_4^{2-}$:$SO_4^{2-}$ ratio, which were performed in triplicates. All experiments are represented as data mean ± standard deviation (s.d.).

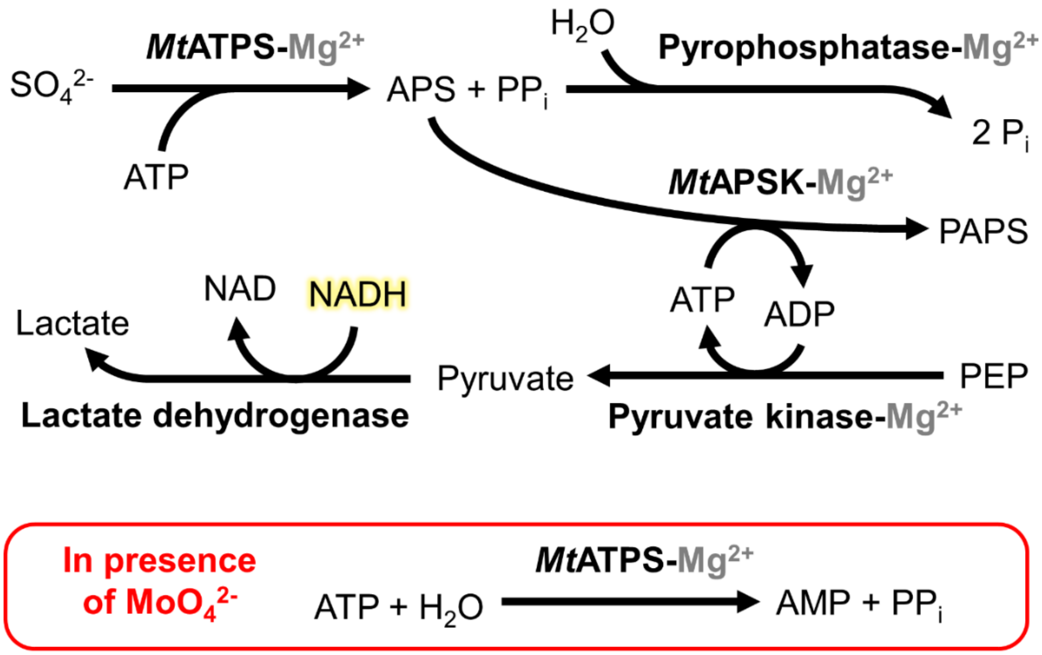


**Supplementary Fig 3. Coupled enzyme assay and $MoO_4^{2-}$ inhibition.** Scheme of the coupled enzyme assay used in this study and impact of $MoO_4^{2-}$ addition. When $MoO_4^{2-}$ binds to the active site of the ATP-sulfurylase ATP is hydrolysed into AMP and $PP_i$ (molybdolysis).

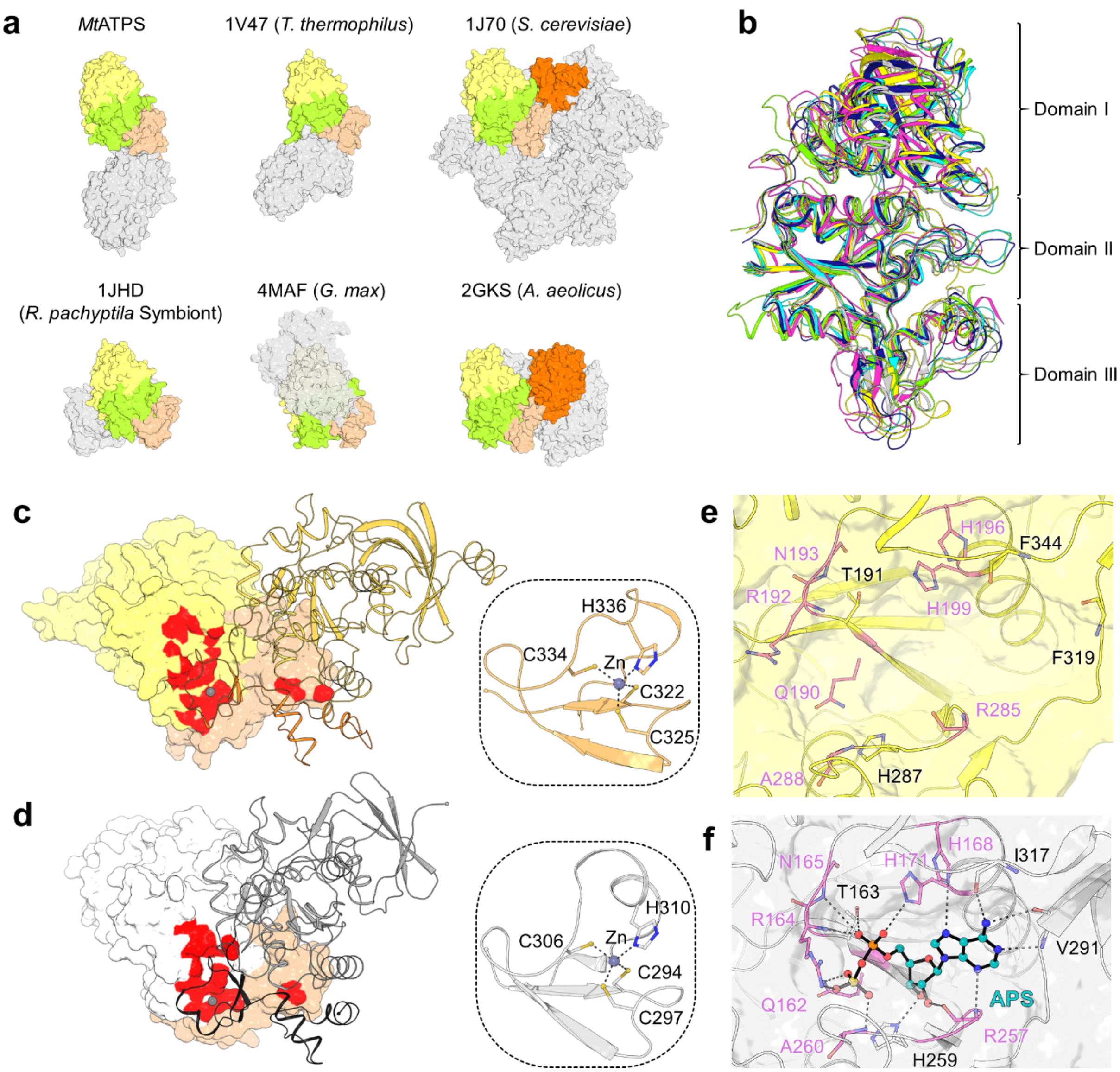


**Supplementary Fig 4. ATP-sulfurylase of *M. thermolithotrophicus* (*Mt*ATPS). a,** Comparison of ATPS in surface representation, coloured by the domain composition I (yellow), II (lemon), III (wheat) and APS kinase (dark orange). The grey surfaces corresponds to the opposing monomer. *Sc*ATPS organizes as homohexamer. **b**, Monomers of *Mt*ATPS (yellow), *Tt*ATPS (grey), *Sc*ATPS (navy blue), *Rrs*ATPS (magenta), *Gm*ATPS (green) and *Aa*ATPS (cyan) are superposed on domain II and shown as cartoons. Abbreviations and rmsd can be found in Supplementary Table 2. **c,d,** Surface area involved in the oligomerization of *Mt*ATPS (c) and *Tt*ATPS (d). One monomer is shown in surface representation and one monomer is displayed in cartoon. The monomer-monomer contacts, established by the domain III of one chain (in orange for *Mt*ATPS and black for *Tt*ATPS), are shown as a red surface. The wheat coloured surface highlights domain III. The framed inlet is a close up of the Zn binding motif and residues coordinating the Zn are drawn as sticks. Carbon, nitrogen and sulfur are coloured as orange/white, blue and yellow, respectively. **e**, **f,** Catalytic site of *Mt*ATPS (apo, e) and *Tt*ATPS with bound APS (f). Elements are coloured as in inlets (c, d) with oxygen and phosphorous in red and orange, respectively. Residues belonging to the canonical motifs of the ATPS are highlighted by pink coloured carbon atoms.

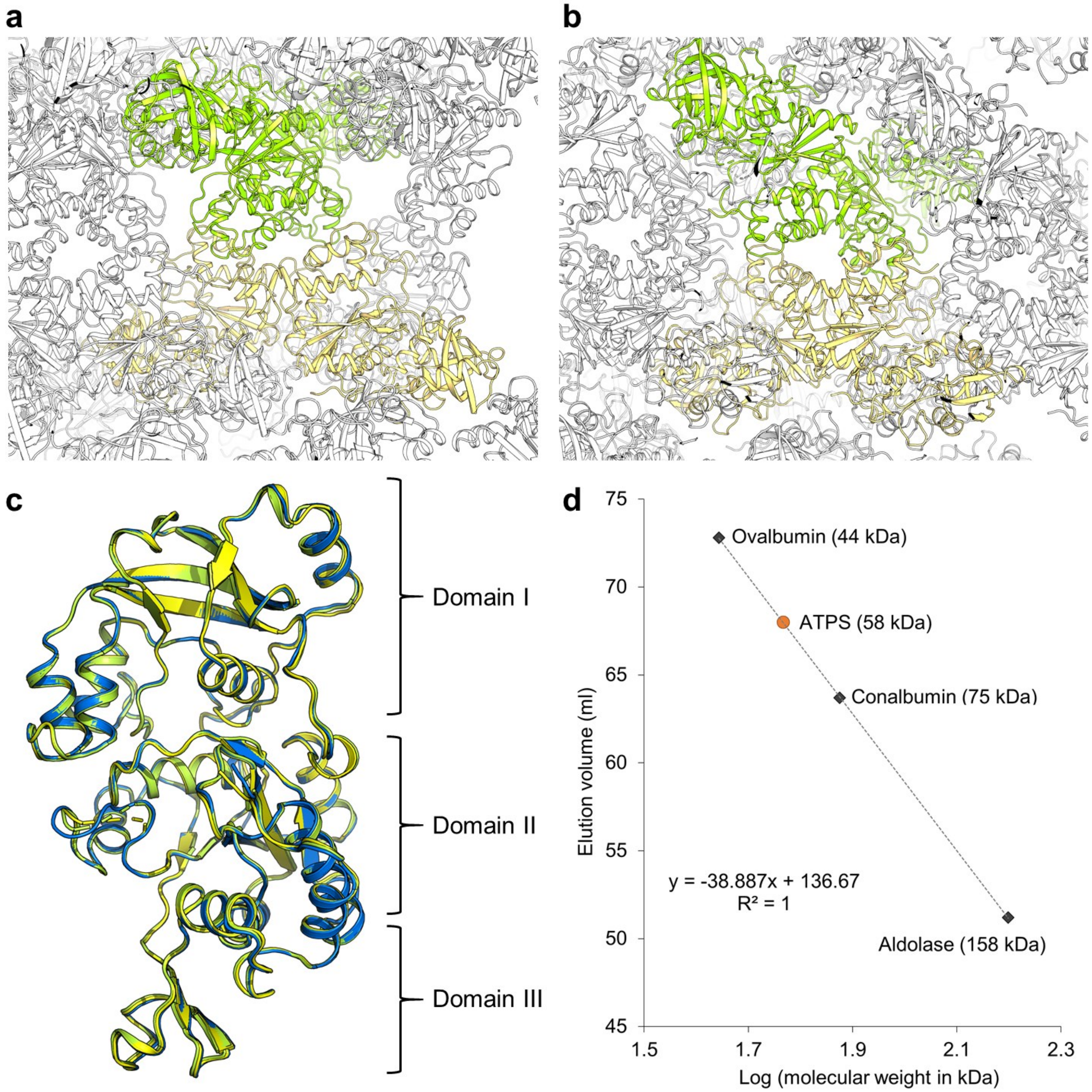


**Supplementary Fig 5. Crystalline packing of *Mt*ATPS suggests a homotetramer. a, b**, Packing of the crystalline form 1 (a) and 2 (b) containing a monomer and a dimer in the asymmetric unit, respectively. All *Mt*ATPS are shown as cartoon with the main dimeric unit coloured in light yellow. The opposite dimer, related by a 2-fold axis symmetry, is coloured in light green and suggest a tetrameric unit, contradicted by the PISA server and gel filtration (see panel d). **c**, The two monomers from *Mt*ATPS form 2 superpose well (rmsd 0.253 Å for 348 Cα, coloured in yellow and green) as well as the monomer from *Mt*ATPS form 1 (rmsd 0.238 Å for 331 Cα, coloured as blue). **d**, Gel filtration profile of *Mt*ATPS and a high molecular weight calibration kit (GE Healthcare). The obtained experimental molecular weight of 58 kDa is smaller than the expected dimeric form (89 kDa calculated from the sequence) but refutes a homotetrameric arrangement.

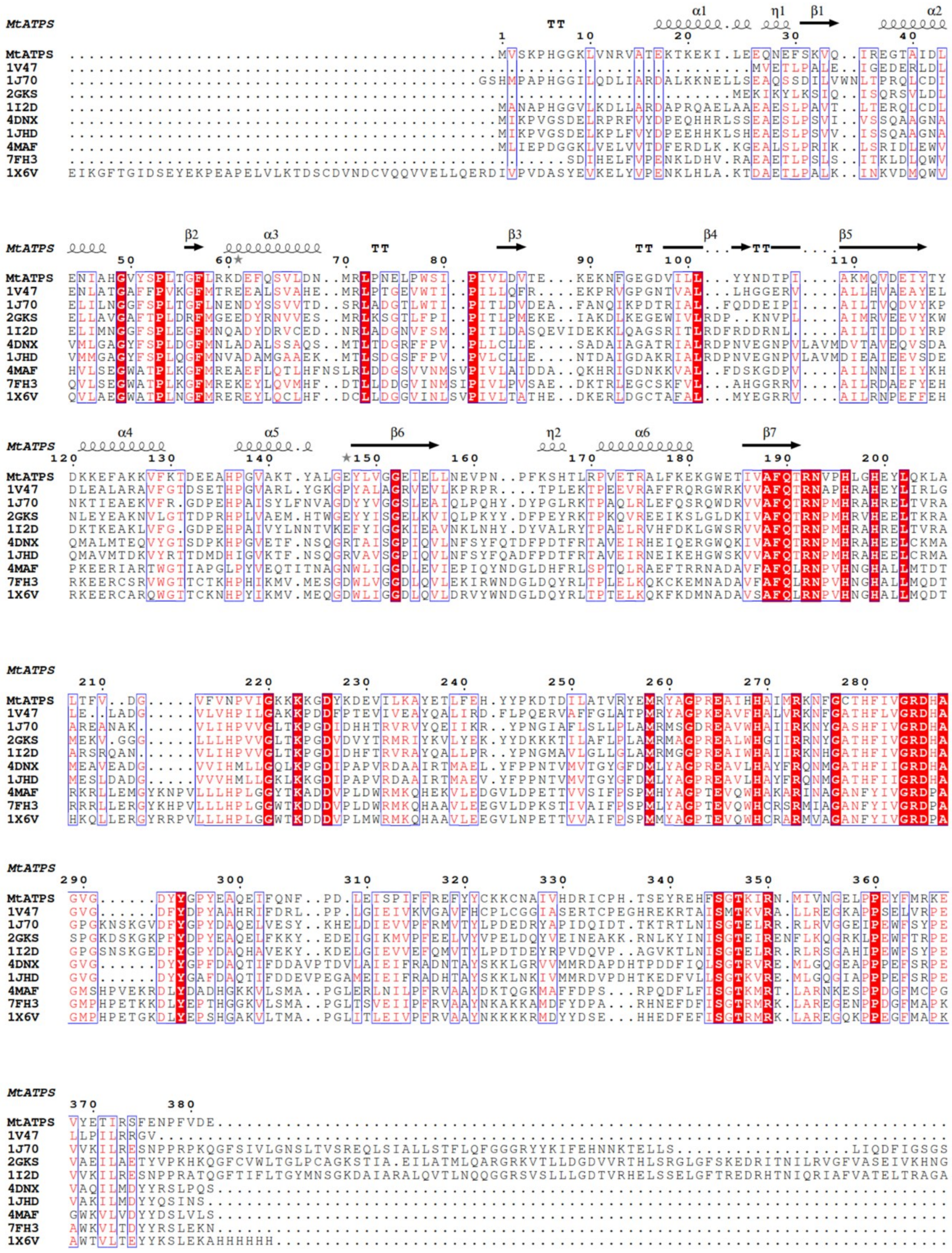

**Supplementary Fig 6. Sequence conservation across *Mt*ATPS with homologues.** Perfectly conserved residues are highlighted with a red background. Sequence alignment was done using MUSCLE[3], secondary structure prediction was performed with ESPript 3.0[4]. For more information regarding the PDB accession numbers, see Supplementary Table 2.

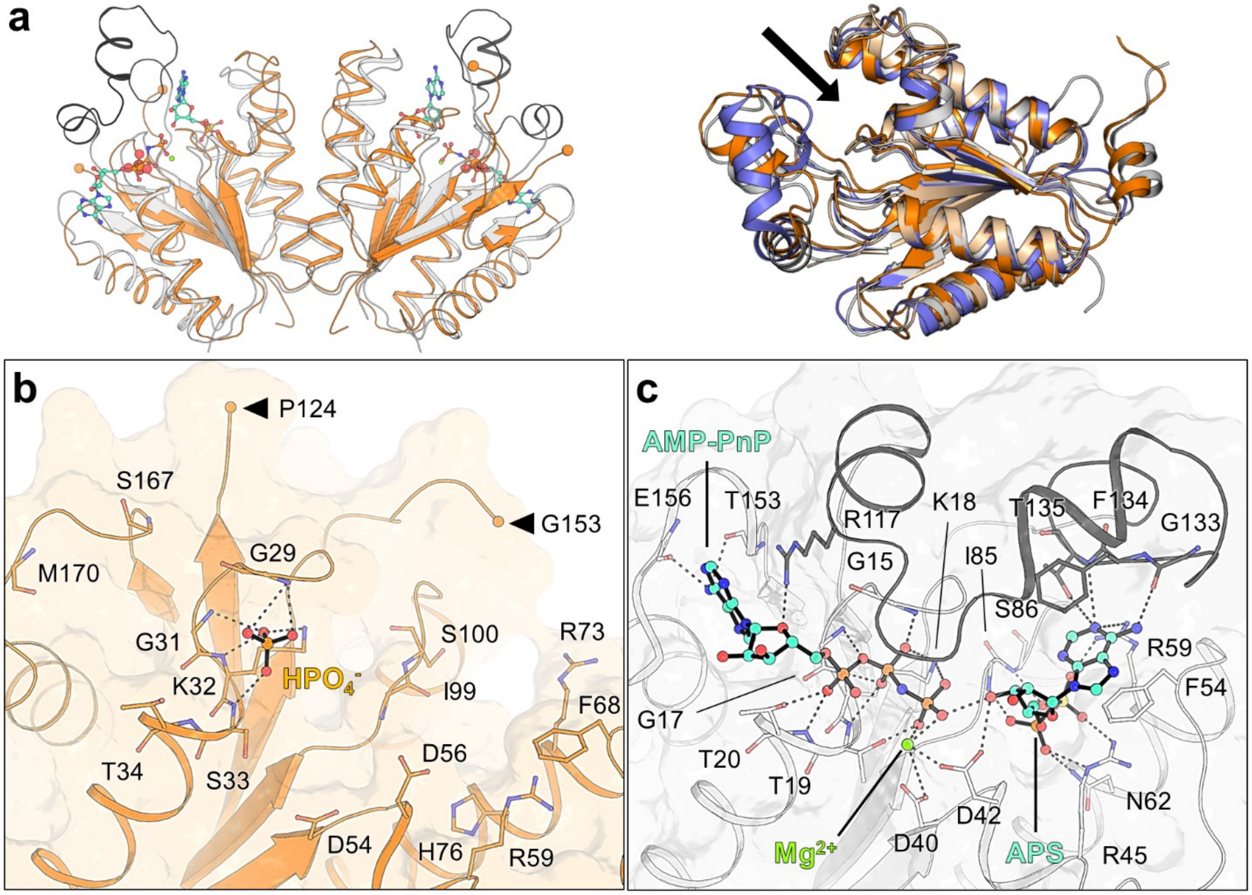


**Supplementary Fig 7. *Mt*APSK belongs to the APSK family**. **a**, Left panel, homodimeric *Mt*APSK apo (orange) superposed to its closest homologue *Synechocystis* sp. PCC 6803 (*Ss*APSK, PDB: 5CB6) in complex with APS and AMP-PnP (white). The ligands are shown in sticks and spheres and the missing part 124-153 in *Mt*APSK (indicated by balls) is highlighted in black in *Ss*APSK. Carbon, nitrogen, oxygen, sulfur and phosphorous are coloured in light orange/white/cyan, blue, red, yellow and orange, respectively. Right panel: superposition of *Mt*APSK (wheat), *At*APSK (orange, PDB: 3UIE), *Ap*APSK (slate, PDB: 2YVU) and *Pc*APSK (white, PDB: 1M7H) on one monomer and shown in cartoon. The active site position is indicated by a black arrow. Abbreviations and rmsd can be found in Supplementary Table 3. **b, c**, Catalytic site of apo *Mt*APSK (b) and *Ss*APSK bound to APS and AMP-PnP (c). Elements are coloured as in (a), left panel. In (b), arrows indicate the missing part 124-153.

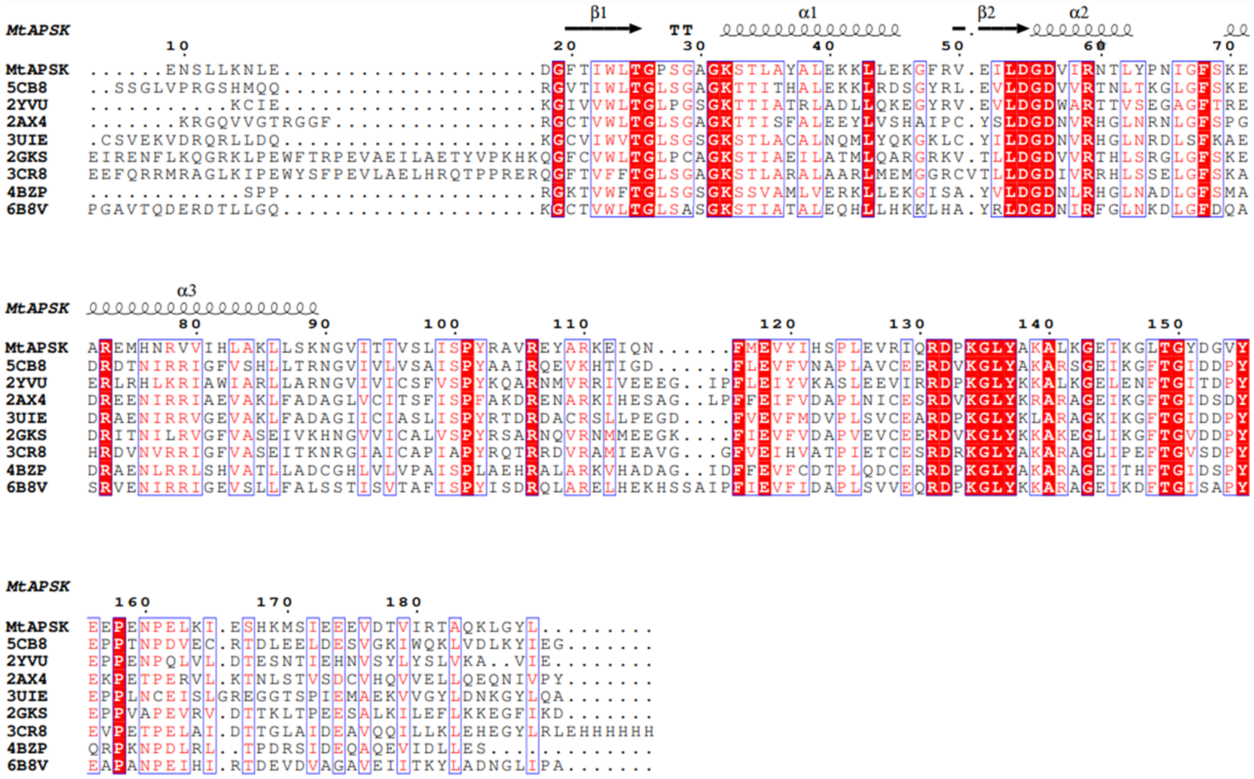


**Supplementary Fig 8. Sequence conservation across *Mt*APSK with other APS-kinases.** Perfectly conserved residues are highlighted with a red background. Sequence alignment was done using MUSCLE[3], secondary structure prediction was performed with ESPript 3.0[4]. For more information regarding the PDB accession numbers, see Supplementary Table 3.

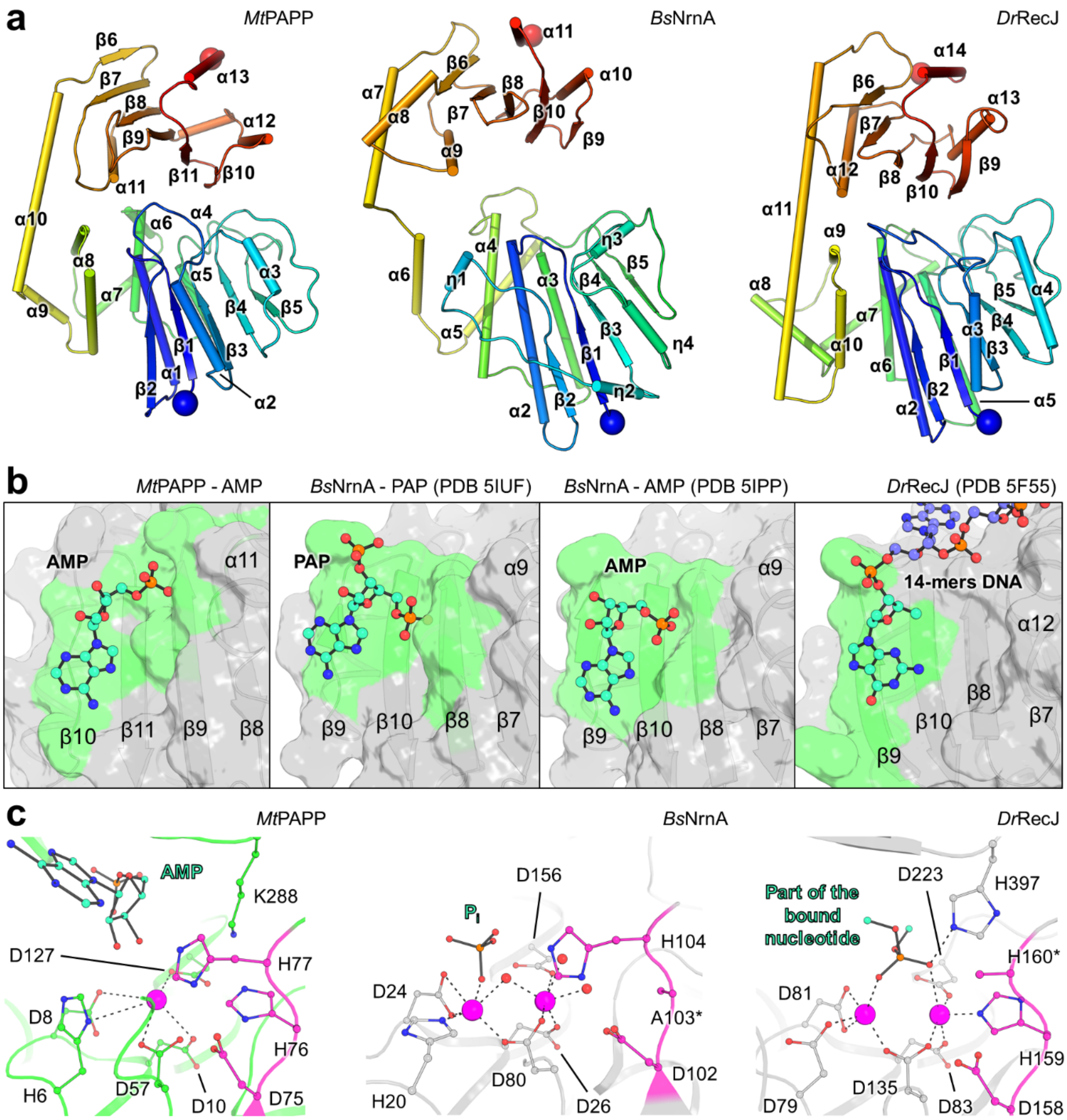


**Supplementary Fig 9. *Mt*PAPP shares similar structural features with exonucleases.** **a**, Folding conservation across the *Mt*PAPP, the NanoRNase A from *Bacillus subtilis* (*Bs*NrnA, PDB: 5IUF) and the recombinase RecJ from *Deinococcus radiodurans* (*Dr*RecJ, PDB: 5F55). For *Bs*NrnA and *Dr*RecJ, the structures only represent the DHH and DHHA1 domains and the secondary structure motifs is renumbered to simplify the comparison with *Mt*PAPP. **b**, Differences in the nucleotide binding between *Mt*PAPP, *Bs*NrnA and *Dr*RecJ. The binding area is represented by a green colour. **c**, Close-up of the $Mn^{2+}$ coordination between *Mt*PAPP, *Bs*NrnA (PDB: 5IZO) and *Dr*RecJ (PDB: 5F55). $Mn^{2+}$ are shown as magenta spheres and the residues coordinating them are highlighted as stick and balls. Carbon, nitrogen, oxygen and phosphorous atoms are coloured respectively in green/grey, blue, red and orange. Residues belonging to the canonical DHH-motif have pink coloured carbon atoms. The structure used for *Bs*NrnA is the His103Ala variant, and the side chain of the His160 has not been modelled in the *Dr*RecJ structure.

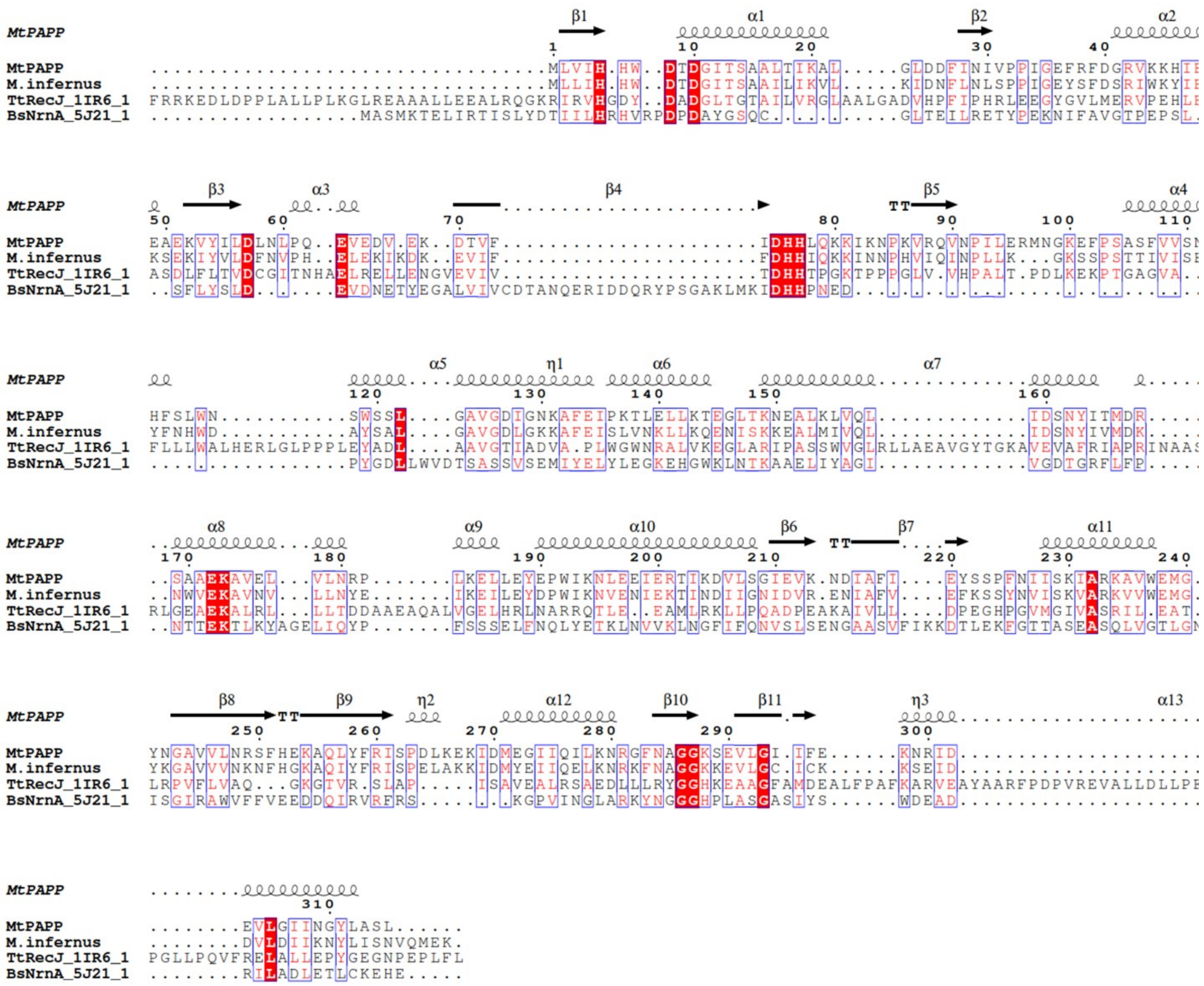


**Supplementary Fig 10. Sequence conservation across *Mt*PAPP, the putative PAPP from *Methanocaldococcus infernus* and structural homologues.** Perfectly conserved residues are highlighted with a red background and a blue box highlights the DHH motif. MtPAPP: *Methanothermococcus thermolithotrophicus* PAP-phosphatase; M.infernus: *Methanocaldococcus infernus* (WP_013099421, putative PAPP); TtRecJ: *Thermus thermophilus* exonuclease RecJ (PDB: 1IR6); BsNrnA_5J21: *Bacillus subtilis* bifunctional oligoribonuclease and PAP-phosphatase NrnA (PDB: 5J21). To allow a correct alignment the “MGSSHHHHHHENLYFQS”-tag from BsNrnA_5J21 was removed. Sequence alignment was done using MUSCLE[3], secondary structure prediction was performed with ESPript 3.0[4].

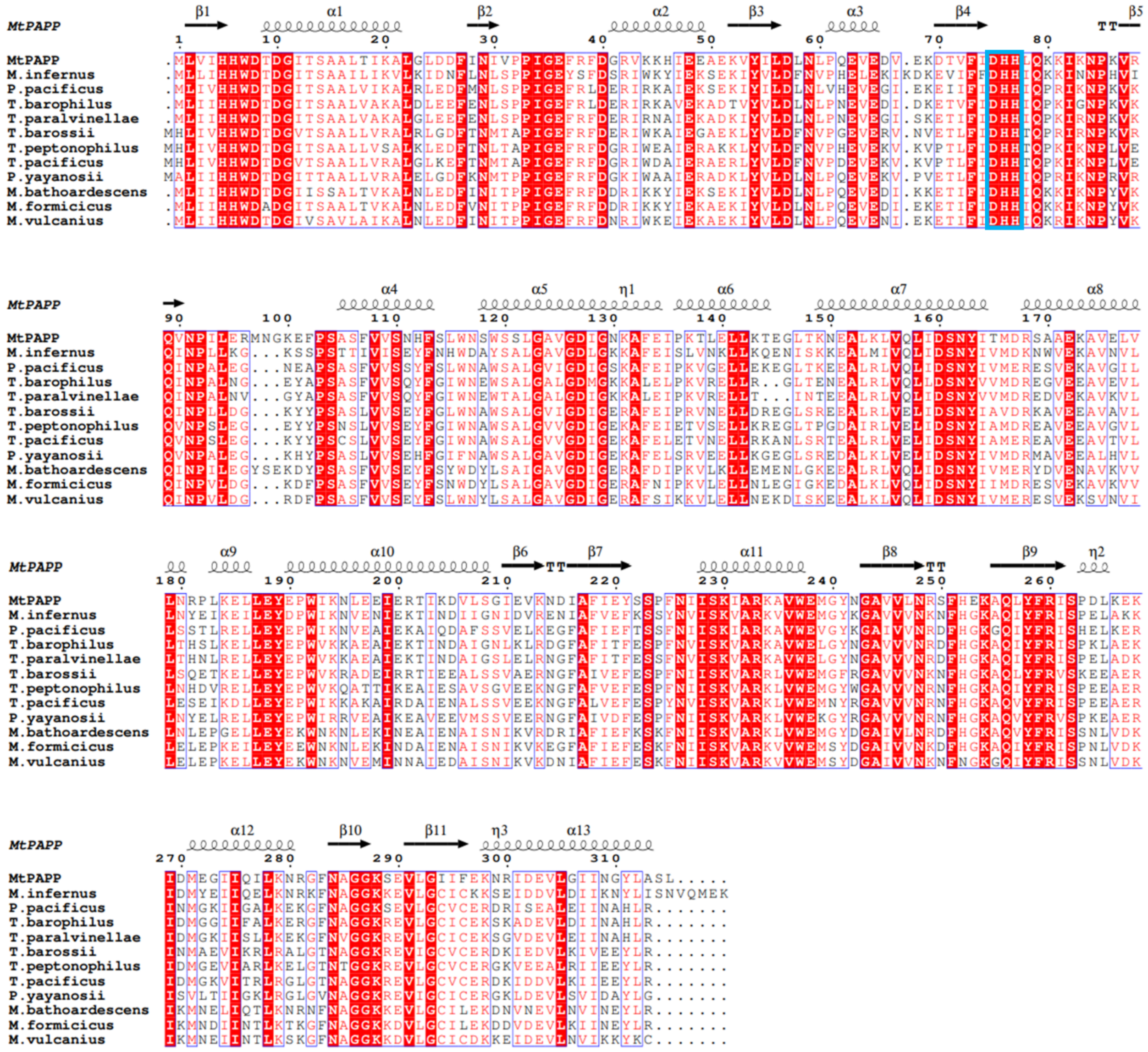


**Supplementary Fig 11. Sequence alignment of *Mt*PAPP with archaeal homologues.** Perfectly conserved residues are highlighted with a red background and a blue box highlights the DHH motif. MtPAPP: *Methanothermococcus thermolithotrophicus* PAP-phosphatase; M.infernus: *Methanocaldococcus infernus* (WP_013099421); P.pacificus: *Palaeococcus pacificus* (WP_048165810); T.barophilus: *Thermococcus barophilus* (WP_013468329); T.paralvinellae: *Thermococcus paralvinellae* (WP_042682556); T.barossii: *Thermococcus barossii* (WP_088865569); T.peptonophilus: *Thermococcus peptonophilus* (WP_062389597); T.pacificus: *Thermococcus pacificus* (WP_088853513); P.yayanosii: *Pyrococcus yayanosii* (WP_048058414); M.bathoardescens: *Methanocaldococcus bathoardescens* (WP_048201137); M.formicicus: *Methanotorris formicicus* (WP_007044583); M.vulcanius: *Methanocaldococcus vulcanius* (WP_012819478). Sequence alignment was done using MUSCLE[3], secondary structure prediction was performed with ESPript 3.0[4].

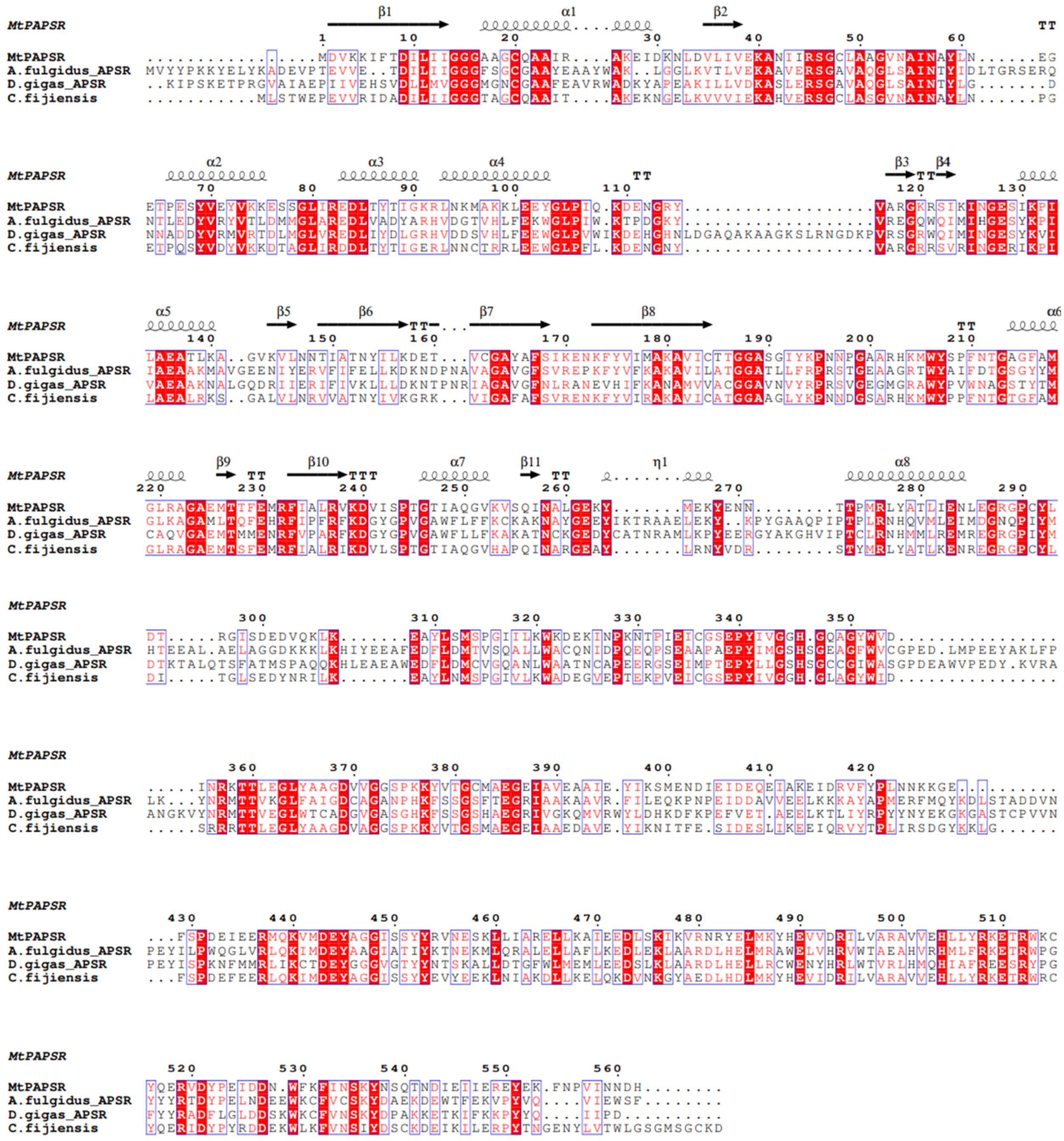


**Supplementary Fig 12. Sequence alignment of dissimilatory APS-reductases with *Mt*PAPSR.** Perfectly conserved residues are highlighted with a red background. MtPAPSR: PAPS-reductase from *Methanothermococcus thermolithotrophicus*, A.fulgidus_APSR: alpha subunit of the APSR from *Archaeoglobus fulgidus* (PDB: 2FJA), D.gigas_APSR: alpha subunit of the APSR from *D. gigas* (PDB: 3GYX), and C.fijiensis: the putative APSR from the bacterium *Caldanaerobius fijiensis* (*Cf*APSR, WP_073344903, closest homologue of *Mt*PAPSR alpha). Sequence alignment was done using MUSCLE[3], secondary structure prediction was performed with ESPript 3.0[4].

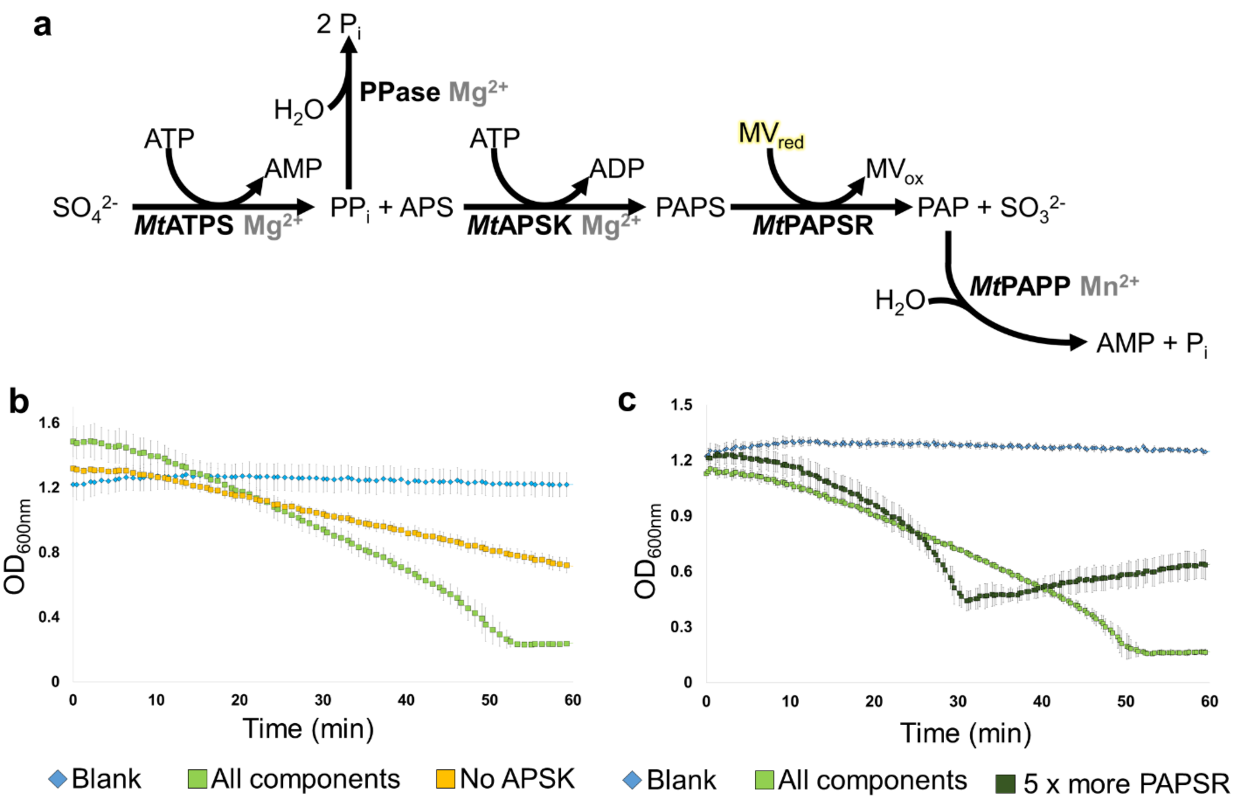


**Supplementary Fig 13. *In vitro* reconstitution of the $SO_4^{2-}$ reduction pathway to measure *Mt*PAPSR activity. a**, Scheme of the coupled enzyme assay used to measure *Mt*PAPSR activity via the oxidation of reduced methyl viologen ($MV_{red}$) at 600 nm under an $N_2$-atmosphere and at 50 °C. The enzymes were recombinantly expressed in *E.coli*, except for the pyrophosphatase, which was commercially obtained. The reaction mix contained all enzymes, metals ($Mn^{2+}$, $Mg^{2+}$), HEPES buffer at pH 7.0 and the substrates $SO_4^{2-}$ and ATP. The reaction was started by the addition of $SO_4^{2-}$. **b, c,** "Blank" corresponds to a solution containing the buffer, the substrates, $MV_{red}$ and dithionite but no enzymes. "All components" contained every compound shown in scheme (a). **b,** Substrate specificity of *Mt*PAPSR. "No *Mt*APSK" corresponds to all components except *Mt*APSK, which prevented the generation of PAPS from APS. With APS as the substrate, a specific enzymatic activity of 0.007 ± 0.001 µmol of oxidized $MV.min^{-1}.mg^{-1}$ of PAPSR was measured, which is only 6.54 % of the activity with the APSK (corresponds to Fig. 4a, +APS -ATPS -APSK). The difference can be attributed to an instability of the added APS. **c**, Impact of different concentrations of *Mt*PAPSR on the activity: A five-fold addition of *Mt*PAPSR resulted in an increase from 0.089 ± 0.007 to 0.251 ± 0.007 µmol of oxidized $MV.min^{-1}.mg^{-1}$ of the *Mt*PAPSR (shown in dark green squares, "5 x more PAPSR"). After 30 minutes the five-fold addition of the *Mt*PAPS-reductase led to aggregation, which is why the $OD_{600nm}$ started to increase after this time point. All experiments were performed in triplicates and represented as data mean ± s.d.

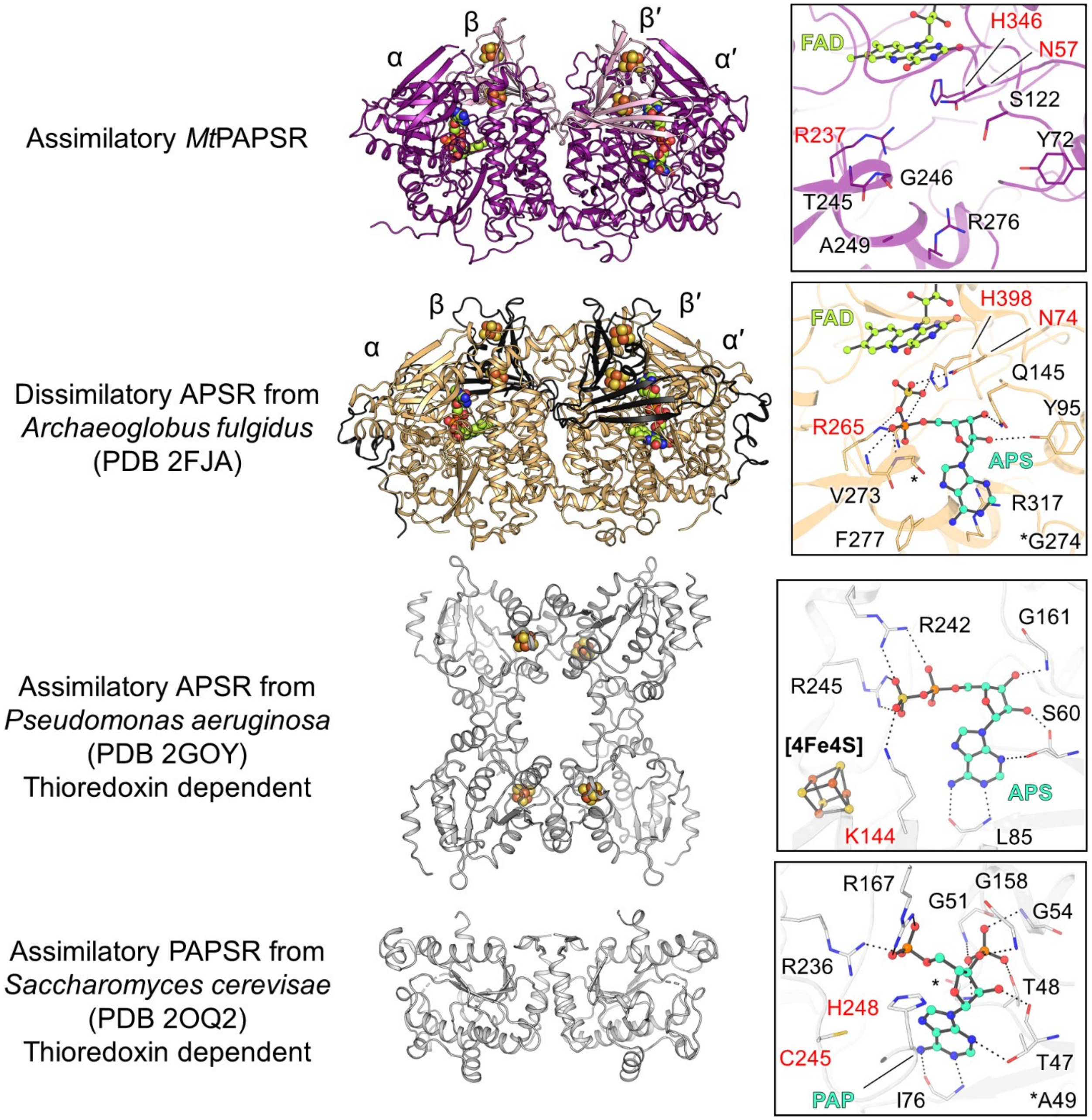


**Supplementary Fig 14. Comparison of a dissimilatory APS-reductase and assimilatory P/APS-reductases with the *Mt*PAPSR**. All structures are shown in cartoon with their (metallo)-cofactors in balls and sticks. Close-ups of the active sites (on the right) with residues important for substrate binding are highlighted as balls and sticks. The dissimilatory APS-reductase from *Archaeoglobus fulgidus* is a heterotetramer, composed of two (αβ)-subunits. Each β-subunit contains two [4Fe-4S]-cluster and each α-subunit one FAD. The presented assimilatory APSR from *Pseudomonas aeruginosa* is a homotetramer. It contains one [4Fe-4S]-cluster per monomer. The assimilatory PAPSR from *Saccharomyces cerevisiae* is homodimeric. The elements oxygen, nitrogen, phosphorous, sulfur and iron are coloured red, blue, orange, yellow and brown. The carbon of the substrate/product is coloured in cyan, and in yellow for the FAD. Catalytic residues are highlighted with red labels.

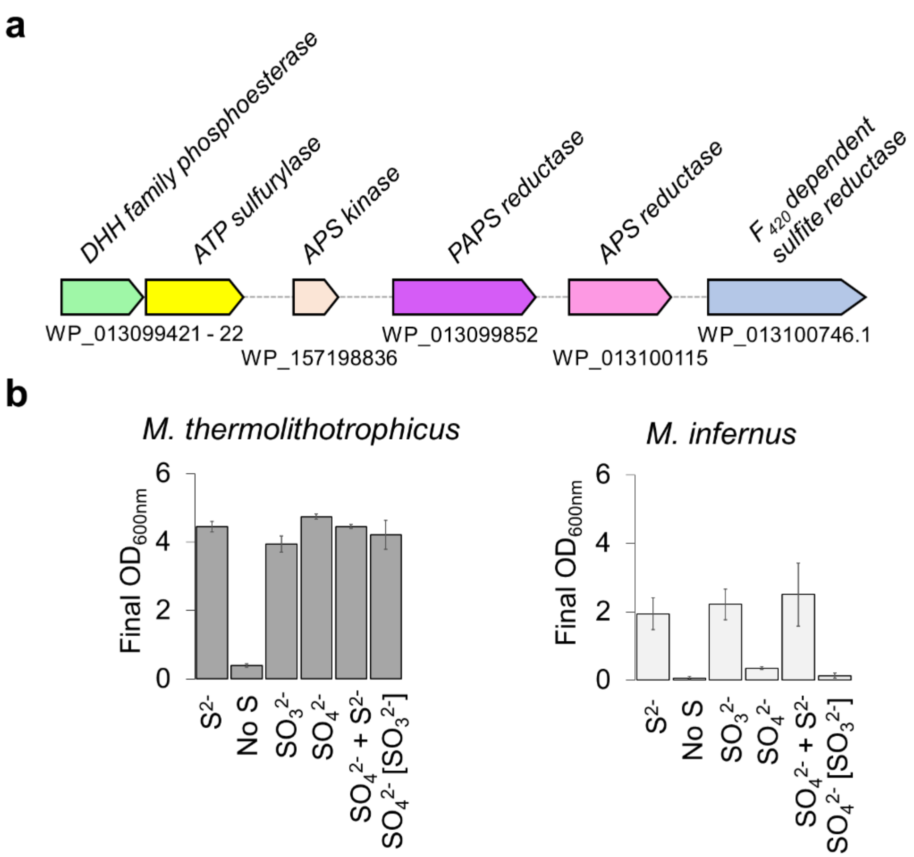


**Supplementary Fig 15. Sulfate-reduction potential in *Methanococcales*. a**, *Methanocaldococcus infernus* has the genomic potential to perform the whole $SO_4^{2-}$ assimilation pathway. WP_013099421 has 59.68 % amino acid sequence identity with *Mt*PAPP, WP_013099422 has 70.60 % sequence identity with *Mt*ATPS, WP_157198836 has 75.44 % sequence identity with *Mt*APSK. The APSR (WP_013100115) and PAPSR (WP_013099852) are similar to the biochemically characterized APSR and PAPSR from *M. jannaschii* (68.64 % and 58.35 % amino acid sequence identity, respectively), which have been shown to reduce APS/PAPS.[5,6] WP_013099852 is not homologous to *Mt*PAPSR but homologous to WP_018154242 a putative PAPS-reductase in *M. thermolithotrophicus*. WP_013100746 has 65.30 % sequence identity to Group-I *Mt*Fsr. **b**, Growth of *M. thermolithotrophicus* and *M. infernus* on 2 mM $Na_2S$, without an additional sulfur source, 2 mM $Na_2SO_3$, 2 mM $Na_2SO_4$ or 2 mM $Na_2SO_4$ with 2 mM $Na_2S$. The $SO_3^{2-}$ in brackets indicates that it was used as the sulfur-substrate for the inoculum. Shown are the maximum optical density measurements ($OD_{600nm}$) of the cultures, in triplicates ± s.d.

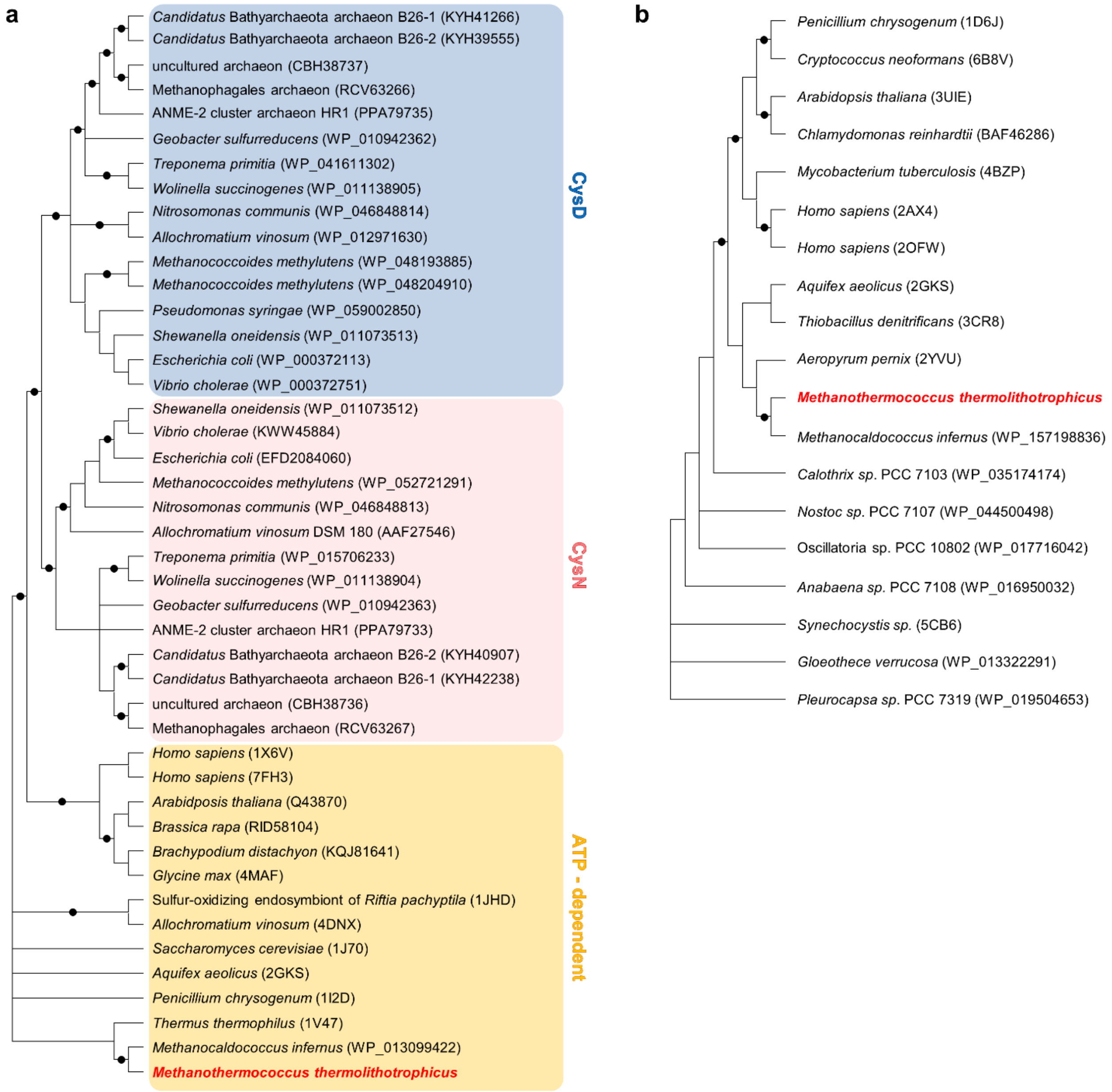


**Supplementary Fig 16**. Phylogenetic analysis of **a,** the heterodimeric sulfate adenylyltransferase (CysDN) with the homo-oligomeric ATP-dependent ATP-sulfurylase (sat) and **b,** APS-kinases. (a) The heterodimeric assimilatory ATP-sulfurylase is composed of a regulatory GTPase subunit CysN (light red) and a catalytic subunit CysD (blue). Sat is involved in both assimilatory and dissimilatory sulfate reduction (light orange). *Mt*ATPS and *Mt*APSK are highlighted in bold red. Bootstrap support values ≥ 90 % are shown as dots on interior nodes.

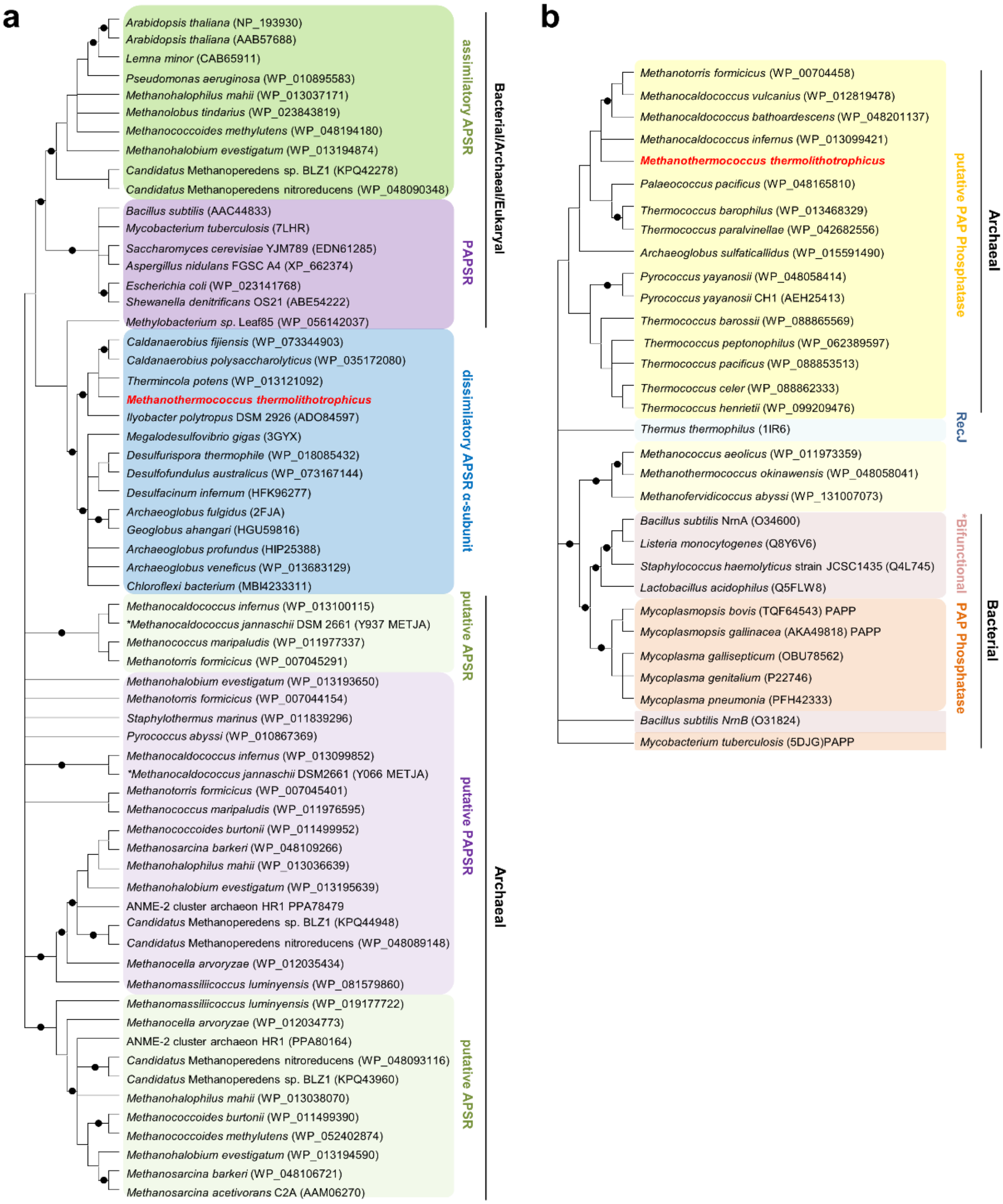


**Supplementary Fig 17**. Phylogenetic analysis of **a,** dissimilatory APS-reductases (α-subunit) and (putative) assimilatory APS/PAPS-reductases, and **b,** PAP-phosphatases with bifunctional oligoribonucleases and PAP-phosphatases, as well as exonucleases (RecJ). *Mt*PAPSR and *Mt*PAPP are highlighted in bold red. Bootstrap support values ≥ 90 % are shown as dots on interior nodes.

| | ATP-sulfurylase (reference WP_018153795) | PAP-phosphatase (reference WP_018153796) | APS-kinase (reference WP_018153797) | PAPS-reductase alpha subunit (reference WP_018153799) |
|---|---|---|---|---|
| **Methanopyrales** | / | / | / | / |
| **Methanococcales** | *Methanotorris formicicus* (WP_048115642.1, 78 %)<br>*Methanocaldococcus bathoardescens* (WP_048201136.1, 79 %)<br>*Methanocaldococcus sp.* SG7 (WP_214399893.1; 73 %)<br>*Methanocaldococcus infernus* (WP_013099422.1; 71 %)<br>*Methanocaldococcus vulcanius* (WP_012819477.1; 77 %)<br>*Methanotorris formicicus* Mc-S-70 (EHP86153.1, 78 %) | *Methanotorris formicicus* (WP_007044583.1; 68 %)<br>*Methanocaldococcus bathoardescens* (WP_048201137.1; 67 %)<br>*Methanocaldococcus* sp. SG7 (WP_214399894.1, 62 %)<br>*Methanocaldococcus infernus* (WP_013099421.1; 60 %)<br>*Methanocaldococcus vulcanius* (WP_012819478.1; 64 %) | *Methanotorris formicicus* (WP_007044585.1; 76 %)<br>*Methanocaldococcus bathoardescens* (WP_048201139.1; 75 %)<br>*Methanocaldococcus sp. SG7* (WP_214399895.1, 76 %)<br>*Methanocaldococcus infernus* (WP_157198836.1; 75 %)<br>*Methanocaldococcus vulcanius* (WP_012819479.1; 76 %) | / |
| **Methanobacteriales** | / | / | / | / |
| **Methanomicrobiales** | *Methanoregulaceae* archaeon (NTV00908.1; 33 %)<br>*Methanomicrobiales* archaeon HGW-Methanomicrobiales-1 (PKL70343.1; 39 %) | / | *Methanoregulaceae archaeon* (RPI39683.1; 47 %)<br>*Methanomicrobiaceae archaeon* (MBN2735040.1; 49 %)<br>*Methanoregula formicica* (WP_015284456.1; 44 %) | / |
| **Methanomassiliicoccales** | / | / | / | / |
| **Methanosarcinales** | *Methanosarcinales* archaeon (RLG37318.1, 57 %)<br>*Methanosarcinales archaeon* (TRZ88670.1, 57 %)<br>*Candidatus* Methanoperedens sp. (NJD52436.1, 57 %)<br>*Methanohalophilus mahii* (WP_013037172.1; 55 %)<br>*Methanohalophilus* sp. RSK (WP_123135802.1; 54 %)<br>+ 23 more | *Methanosarcinales* archaeon (RLG38197.1; 28 %)<br>*Methanosarcinales* archaeon (MCD4846224.1, 32 %)<br>*Methanohalobium evestigatum* (WP_013194595.1; 33 %) | *Methanosarcinales* archaeon (RLG37639.1; 53 %)<br>*Methanosarcinales* archaeon (MCD4846223.1; 50 %)<br>*Methanohalobium evestigatum* (WP_013194596.1; 51 %)<br>*Methanococcoides orientis* (WP_233084240.1; 49 %)<br>*Methanohalophilus profundi* (WP_129597714.1; 48 %)<br>*Methanohalophilus portucalensis* (WP_072358461.1; 50 %)<br>+ 25 more | *Methanosarcinales* archaeon (RLG30695.1; 34 %)<br>*Methermicoccus shengliensis* (WP_052353262.1; 40 %)<br>*Candidatus* Methanoperedenaceae archaeon GB37 (CAD7771153.1; 36 %)<br>*Candidatus* Methanoperedenaceae archaeon GB50 (CAD7770084.1; 36 %) |
| **Methanofastidiosa** | / | / | / | / |

**Supplementary Fig 18. $SO_4^{2-}$-reduction associated genes across the seven orders of methanogens.** The protein sequences from the biochemically and structurally characterized enzymes from *M. thermolithotrophicus* were used as reference. The NCBI accession numbers of the proteins (left) as well as the amino acid sequence identity (in %) in comparison to *M. thermolithotrophicus* enzymes are shown in brackets. Homologues below 28 % sequence identity or with a coverage below 85 % are not shown.

**Supplementary Table 1.** X-ray analysis statistics for the presented structures.

| | ***Mt*ATPS form 1** | ***Mt*ATPS form 2** | ***Mt*APSK** | ***Mt*PAPP** | ***Mt*PAPSR Fe K edge** | ***Mt*PAPSR** |
|---|---|---|---|---|---|---|
| **Data collection** | | | | | | |
| Synchrotron source | SOLEIL, Proxima-I | SOLEIL Proxima-I | SLS, X06DA | SLS, X06DA | PETRAIII, P11 | SLS, X06DA |
| Wavelength (Å) | 1.00000 | 1.03320 | 1.00003 | 1.64566 | 1.73646 | 0.97625 |
| Space group | *I*222 | *C*2 | *P*$2_1$ | *I*4 | *P*$2_1$ | *P*$2_1$ |
| Resolution (Å) | 78.79 – 1.97 (2.22– 1.97) | 65.94-2.10 (2.16-2.10) | 47.74– 1.77 (1.81– 1.77) | 126.38 – 3.10 (3.27 – 3.10) | 85.51 – 1.89 (2.17 – 1.89) | 86.04 – 1.45 (1.64 – 1.45) |
| Cell dimensions | | | | | | |
| a, b, c (Å) | 55.70, 154.46, 157.57 | 185.28, 54.52, 85.43 | 51.54, 176.18, 51.61 | 174.05, 174.05, 183.80 | 63.53, 122.82, 88.53 | 63.71, 123.65, 88.84 |
| α, β, γ (°) | 90, 90, 90 | 90, 95.91, 90 | 90, 105.78, 90 | 90, 90, 90 | 90, 105.01, 90 | 90, 104.44, 90 |
| $R_{merge}$(%)[a] | 15.1 ( 174.0) | 13.5 (114.9) | 4.4 (92.8) | 19.7 (248.3) | 20.0 (169.8) | 7.9 (105.6) |
| $R_{pim}$ (%)[a] | 4.2 (48.3) | 5.5 (47.5) | 1.8 (39.8) | 5.6 (68.0) | 6.9 (62.5) | 3.3 (42.5) |
| $CC_{1/2}$[a] | 0.998 (0.641) | 0.997 (0.57) | 1.0 (0.685) | 0.998 (0.477) | 0.998 (0.6) | 0.997 ( 0.567) |
| $I/\sigma_I$[a] | 12.7 (1.6) | 10.2 (1.6) | 22.7 (1.7) | 12.3 (1.1) | 12.0 (1.8) | 11.9 (1.7) |
| Spherical completeness[a] | 58.2 (9.9) | 82.3 (51.2) | 80.0 (64.5) | 92.1 (100.0) | 55.7 (8.3) | 62.3 (10.2) |
| Ellipsoidal completeness[a] | 93.2 (62.3) | 95.6 (98.3) | 83.1 (96.3) | / | 91.8 (59.4) | 95.0 (73.8) |
| Redundancy[a] | 13.7 (13.9) | 6.9 (6.6) | 6.8 (6.2) | 13.3 (14.3) | 18.2 (14.4) | 6.8 (7.1) |
| Nr. unique reflections[a] | 28,197 (1,410) | 41,069 (2,053) | 68,557 (3,427) | 45,681 (7,240) | 58,277 (2,915) | 146,296 (7,316) |
| **Refinement** | | | | | | |
| Resolution (Å) | 49.73 – 1.97 | 52.28 – 2.10 | 44.04 – 1.77 | 57.79 – 3.10 | | 1.45 |
| Number of reflections | 28,185 | 41,057 | 68,551 | 45,643 | | 14,6283 |
| $R_{work}/R_{free}$[b] (%) | 18.85/21.79 | 19.86/23.94 | 15.40/18.09 | 18.57/21.96 | | 15.14/17.80 |
| Number of atoms | | | | | | |
| Protein | 3,155 | 6,205 | 4,753 | 15,234 | | 10368 |
| Ligands/ions | 49 | 71 | 88 | 183 | | 223 |
| Solvent | 208 | 354 | 606 | 0 | | 879 |
| Mean B-value ($Å^2$) | 42.81 | 39.09 | 37.12 | 94.96 | | 32.18 |
| Molprobity clash score, all atoms | 2.99 | 2.09 | 3.38 | 4.67 | | 3.34 |
| Ramachandran plot | | | | | | |
| Favoured regions (%) | 97.11 | 97.45 | 99.30 | 96.92 | | 97.75 |
| Outlier regions (%) | 0 | 0 | 0 | 0 | | 0 |
| rmsd[c] bond lengths (Å) | 0.011 | 0.005 | 0.007 | 0.004 | | 0.007 |
| rmsd[c] bond angles (°) | 1.138 | 0.823 | 0.935 | 0.613 | | 0.917 |
| **PDB ID code** | 8A8G | 8A8D | 8A8H | 8A8K | | 8A8O |

[a] Values relative to the highest resolution shell are within parentheses. [b] $R_{free}$ was calculated as the $R_{work}$ for 5 % of the reflections that were not included in the refinement. [c] rmsd, root mean square deviation.

**Supplementary Table 2. Sequence and structural alignment of *Mt*ATPS.** Sequence alignment was performed using PyMOL version 2.2.0 (Schrödinger, LLC).

| Name of the organisms | Abbreviation | PDB code | Alignment on Domain II Rmsd in Å (aligned Cα) | Overall alignment Rmsd in Å (aligned Cα) |
|---|---|---|---|---|
| *Methanothermococcus thermolithotrophicus* | *Mt*ATPS | | | |
| *Thermus thermophilus* | *Tt*ATPS | 1V47 | 1.07 (121) | 1.78 (296) |
| *Saccharomyces cerevisiae* | *Sc*ATPS | 1J70 | 1.06 (112) | 1.61 (266) |
| *Aquifex aeolicus* | *Aa*ATPS | 2GKS | 1.25 (125) | 1.55 (272) |
| *Penicillium chrysogenum* | *Pc*ATPS | 1I2D | 1.12 (112) | 1.54 (279) |
| *Allochromatium vinosum* | *Av*ATPS | 4DNX | 0.83 (116) | 1.44 (257) |
| *Ryftia pachyptila symbiont* | *Rrs*ATPS | 1JHD | 0.92 (124) | 1.42 (260) |
| *Glycine max* | *Gm*ATPS | 4MAF | 1.27 (110) | 2.33 (315) |
| *Homo sapiens* | *Hs*ATPS1 | 1X6V | 1.25 (105) | 1.98 (252) |
| *Homo sapiens* | *Hs*ATPS2 | 7FH3 | 1.10 (105) | 1.83 (261) |

**Supplementary Table 3. Sequence and structural alignment of *Mt*APSK.** Sequence alignment was performed using PyMOL version 2.2.0 (Schrödinger, LLC).

| Name of the organisms | Abbreviation | PDB code | Overall alignment Rmsd in Å (aligned Cα) |
|---|---|---|---|
| *Methanothermococcus thermolithotrophicus* | *Mt*APSK | | |
| *Synechocystis sp.* PCC 6803 | *Ss*APSK | 5CB8 | 1.33 (129) |
| *Arabidopsis thaliana* | *At*APSK | 3UIE | 1.59 (104) |
| *Aeropyrum pernix* | *Ap*APSK | 2YVU | 1.18 (117) |
| *Penicillium chrysogenum* | *Pc*APSK | 1M7H | 1.00 (120) |
| *Homo sapiens* | *Hs*APSK1 | 2OFW | 1.13 (111) |
| *Homo sapiens* | *Hs*APSK2 | 2AX4 | 1.25 (123) |
| *Aquifex aeolicus* | *Aa*APSK | 2GKS | 1.77 (124) |
| *Thiobacillus denitrificans* | *Td*APSK | 3CR8 | 1.14 (116) |
| *Mycobacterium tuberculosis* | *Mt*APSK | 4BZP | 1.13 (111) |
| *Cryptococcus neoformans* | *Cn*APSK | 6B8V | 0.81 (111) |

### Constructs and gene codon optimisation.

**ATP-sulfurylase sequence from *M. thermolithotrophicus***
**(NCBI Accession number: WP_018153795.1)**
MVSKPHGGKLVNRVATEKTKEKILEEQNEFSKVQIREGTAIDLENIAHGVYSPLTGFLRKDEFQSVLDNMR
LPNELPWSIPIVLDVTEKEKNFGEGDVILLYYNDTPIAKMQVDEIYTYDKKEFAKKVFKTDEEAHPGVAKT
YALGEYLVGGEIELLNEVPNPFKSHTLRPVETRALFKEKGWETIVAFQTRNVPHLGHEYLQKLALTFVDGV
FVNPVIGKKKKGDYKDEVILKAYETLFEHYYPKDTDILATVRYEMRYAGPREAIHHAIMRKNFGCTHFIVG
RDHAGVGDYYGPYEAQEIFQNFPDLEISPIFFREFYYCKKCNAIVHDRICPHTSEYREHFSGTKIRNMIVNGE
LPPEYFMRKEVYETIRSFENPFVDE

**Codon optimized *Mt*ATPS sequence cloned into pET-28a(+):**

**CATATG**GTTAGCAAGCCGCACGGTGGCAAACTGGTGAATCGTGTGGCGACCGAGAAGACCAAGGAGA
AGATCCTGGAAGAACAGAACGAATTCAGCAAGGTGCAGATCCGTGAGGGTACCGCGATCGACCTGGA
AAACATTGCGCATGGTGTGTACAGCCCGCTGACCGGCTTCCTGCGTAAAGACGAGTTTCAAAGCGTTC
TGGATAACATGCGTCTGCCGAACGAACTGCCGTGGAGCATCCCGATTGTGCTGGATGTTACCGAGAAG
GAGAAGAACTTTGGCGAGGGCGACGTGATTCTGCTGTACTATAACGATACCCCGATCGCGAAGATGCA
GGTTGACGAGATTTACACCTATGATAAGAAAGAATTCGCGAAGAAAGTGTTTAAGACCGACGAGGAA
GCGCACCCGGGTGTTGCGAAAACCTACGCGCTGGGCGAGTATCTGGTGGGTGGCGAGATCGAACTGCT
GAACGAAGTTCCGAACCCGTTCAAGAGCCACACCCTGCGTCCGGTTGAAACCCGTGCGCTGTTCAAGG
AGAAAGGTTGGGAAACCATTGTGGCGTTTCAGACCCGTAACGTTCCGCACCTGGGTCACGAATACCTG
CAAAAACTGGCGCTGACCTTCGTGGATGGCGTGTTTGTTAACCCGGTTATCGGTAAGAAAAAGAAAGG
CGACTACAAGGATGAAGTGATTCTGAAAGCGTACGAAACCCTGTTCGAACACTACTATCCGAAGGACA
CCGATATCCTGGCGACCGTTCGTTACGAGATGCGTTATGCGGGTCCGCGTGAAGCGATCCACCATGCG
ATTATGCGTAAAAACTTCGGTTGCACCCACTTTATTGTGGGTCGTGACCACGCGGGTGTTGGTGATTAC
TATGGCCCGTATGAGGCGCAGGAAATTTTCCAAAACTTTCCGGACCTGGAGATCAGCCCGATTTTCTTT
CGTGAATTCTACTATTGCAAGAAATGCAACGCGATCGTGCACGATCGTATTTGCCCGCACACCAGCGA
GTACCGTGAACACTTTAGCGGTACCAAAATCCGTAACATGATTGTTAACGGCGAGCTGCCGCCGGAAT
ATTTTATGCGTAAGGAAGTTTATGAGACCATCCGTAGCTTTGAGAACCCGTTTGTTGATGAG**TGA**GGA
TCC

**Restriction sites (NdeI and BamHI)**

**APS-kinase sequence from *M. thermolithotrophicus***
**(NCBI Accession number: WP_018153797.1)**
MSEELNNGENSLLKNLEDGFTIWLTGPSGAGKSTLAYALEKKLLEKGFRVEILDGDVIRNTLYPNIGFSKEA
REMHNRVVIHLAKLLSKNGVITIVSLISPYRAVREYARKEIQNFMEVYIHSPLEVRIQRDPKGLYAKALKGEI
KGLTGYDGVYEEPENPELKIESHKMSIEEEVDTVIRTAQKLGYL

**Codon optimized *Mt*APSK sequence cloned into pET-28a(+):**

**CATATG**AGCGAGGAACTGAACAACGGCGAAAACAGCCTGCTGAAGAACCTGGAGGACGGCTTCACCA
TTTGGCTGACCGGTCCGAGCGGTGCGGGCAAGAGCACCCTGGCGTACGCGCTGGAAAAGAAACTGCT
GGAGAAAGGCTTCCGTGTGGAAATCCTGGACGGTGATGTTATTCGTAACACCCTGTATCCGAACATTG
GCTTTAGCAAGGAAGCGCGTGAGATGCACAACCGTGTGGTTATCCACCTGGCGAAGCTGCTGAGCAAA
AACGGTGTGATCACCATTGTTAGCCTGATCAGCCCGTACCGTGCGGTGCGTGAATATGCGCGTAAAGA
GATCCAGAACTTTATGGAAGTGTACATTCACAGCCCGCTGGAAGTGCGTATCCAACGTGACCCGAAGG
GCCTGTATGCGAAGGCGCTGAAAGGTGAAATTAAAGGTCTGACCGGCTACGATGGTGTTTATGAGGAA

CCGGAAAACCCGGAGCTGAAGATCGAGAGCCACAAAATGAGCATTGAGGAAGAGGTGGATACCGTTA
TCCGTACCGCGCAGAAACTGGGTTACCTG**TGA**GGATCC

**Restriction sites (NdeI and BamHI)**

**PAP-phosphatase sequence from *M. thermolithotrophicus***

**(NCBI Accession number: WP_018153796.1)**
MLVIHHWDTDGITSAALTIKALGLDDFINIVPPIGEFRFDGRVKKHIEEAEKVYILDLNLPQEVEDVEKDTVF
IDHHLQKKIKNPKVRQVNPILERMNGKEFPSASFVVSNHFSLWNSWSSLGAVGDIGNKAFEIPKTLELLKTE
GLTKNEALKLVQLIDSNYITMDRSAAEKAVELVLNRPLKELLEYEPWIKNLEEIERTIKDVLSGIEVKNDIAF
IEYSSPFNIISKIARKAVWEMGYNGAVVLNRSFHEKAQLYFRISPDLKEKIDMEGIIQILKNRGFNAGGKSEV
LGIIFEKNR IDEVLGIINGYLASL

**Codon optimized *Mt*PAPP sequence cloned into pET-28a(+):**

CATATGCTGGTGATTCACCACTGGGACACCGATGGTATCACCAGCGCGGCGCTGACCATTAAAGCGCT
GGGTCTGGACGATTTCATCAACATTGTTCCGCCGATCGGCGAGTTCCGTTTTGACGGTCGTGTGAAGAA
ACACATCGAGGAAGCGGAAAAAGTTTACATTCTGGATCTGAACCTGCCGCAGGAAGTGGAAGACGTT
GAGAAGGATACCGTGTTTATCGACCACCACCTGCAGAAGAAAATTAAGAACCCGAAAGTGCGTCAAG
TTAACCCGATCCTGGAGCGTATGAACGGCAAAGAGTTCCCGAGCGCGAGCTTTGTGGTTAGCAACCAC
TTCAGCCTGTGGAACAGCTGGAGCAGCCTGGGTGCGGTGGGTGATATCGGTAACAAGGCGTTTGAGAT
TCCGAAAACCCTGGAGCTGCTGAAGACCGAAGGTCTGACCAAGAACGAAGCGCTGAAACTGGTTCAA
CTGATCGACAGCAACTACATTACGATGGACCGTAGCGCGGCGGAGAAGGCGGTGGAACTGGTTCTGA
ACCGTCCGCTGAAAGAGCTGCTGGAGTATGAACCGTGGATTAAGAACCTGGAGGAAATCGAACGTAC
CATTAAAGACGTGCTGAGCGGCATCGAGGTTAAGAACGATATCGCGTTCATTGAATACAGCAGCCCGT
TTAACATCATTAGCAAGATTGCGCGTAAAGCGGTTTGGGAGATGGGCTACAACGGTGCGGTGGTTCTG
AACCGTAGCTTCCACGAAAAAGCGCAGCTGTATTTTCGTATCAGCCCGGACCTGAAGGAGAAAATTGA
TATGGAAGGCATCATTCAAATCCTGAAAAACCGTGGTTTCAACGCGGGTGGCAAGAGCGAAGTGCTG
GGTATCATTTTTGAGAAGAACCGTATCGACGAAGTTCTGGGCATCATTAACGGTTATCTGGCGAGCCT
GT**TGA**GGATCC

**Restriction sites (NdeI and BamHI)**

**PAPS-reductase subunit sequences from *M. thermolithotrophicus***

***Mt*PAPSR subunit alpha sequence**
**(NCBI Accession number: WP_018153799.1)**
MDVKKIFTDILIIGGGAAGCQAAIRAKEIDKNLDVLIVEKANIIRSGCLAAGVNAINAYLNEGETPESYVEYV
KKESSGLIREDLTYTIGKRLNKMAKKLEEYGLPIQKDENGRYVARGKRSIKINGESIKPILAEATLKAGVKV
LNNTIATNYILKDETVCGAYAFSIKENKFYVIMAKAVICTTGGASGIYKPNNPGAARHKMWYSPFNTGAGF
AMGLRAGAEMTTFEMRFIALRVKDVISPTGTIAQGVKVSQINALGEKYMEKYENNTTPMRLYATLIENLEG
RGPCYLDTRGISDEDVQKLKEAYLSMSPGIILKWKDEKINPKNTPIEICGSEPYIVGGHGQAGYWVDINRKT
TLEGLYAAGDVVGGSPKKYVTGCMAEGEIAVEAAIEYIKSMENDIEIDEQEIAKEIDRVFYPLNNKKGEFSP
DEIEERMQKVMDEYAGGISSYYRVNESKLLIARELLKAIEEDLSKIKVRNRYELMKYHEVVDRILVARAVV
EHLLYRKETRWKCYQERVDYPEIDDNWFKFINSKYNSQTNDIEIIEREYEKFNPVINNDH

***Mt*PAPSR subunit beta sequence**
**(NCBI Accession number: WP_018153800.1)**
MTIRIIEEICIGCGLCTKVCPGNLLYQREDGKSEIMDKRDCWDCAACVKECPVNAIEMYLQPEIGGRGSTLK
AKKTDDSIVWIITDNNGEEEVIEVKNKKTFDM

**Codon optimized PAPS-reductase sequences cloned into pET-28a(+):**

**CCATGG*ATG***GATGTTAAGAAGATATTCACAGATATACTCATAATAGGTGGTGGTGCAGCAGGTTGCCA
GGCAGCAATAAGGGCAAAGGAGATAGATAAGAACCTCGATGTTCTCATAGTTGAGAAGGCAAACATA
ATAAGGTCAGGTTGCCTCGCAGCAGGTGTTAACGCAATAAACGCATACCTCAACGAGGGTGAGACAC
CCGAGTCATACGTTGAGTACGTTAAGAAGGAGTCATCAGGTCTCATAAGGGAGGATCTCACATACACA
ATAGGTAAGAGGCTCAACAAGATGGCAAAGAAGCTCGAGGAGTACGGTCTCCCCATACAGAAGGATG
AGAACGGTAGGTACGTTGCAAGGGGTAAGAGGTCAATAAAGATAAACGGTGAGTCAATAAAGCCCAT
ACTCGCAGAGGCAACACTCAAGGCAGGTGTTAAGGTTCTCAACAACACAATAGCAACAAACTACATA
CTCAAGGATGAGACAGTTTGCGGTGCATACGCATTCTCAATAAAGGAGAACAAGTTCTACGTTATAAT
GGCAAAGGCAGTTATATGCACAACAGGTGGTGCATCAGGTATATACAAGCCCAACAACCCCGGTGCA
GCAAGGCACAAGATGTGGTACTCACCCTTCAACACAGGTGCAGGTTTCGCAATGGGTCTCAGGGCAGG
TGCAGAGATGACAACATTCGAGATGAGGTTCATAGCACTCAGGGTTAAGGATGTTATATCACCCACAG
GTACAATAGCACAGGGTGTTAAGGTTTCACAGATAAACGCACTCGGTGAGAAGTACATGGAGAAGTA
CGAGAACAACACAACACCCATGAGGCTCTACGCAACACTCATAGAGAACCTCGAGGGTAGGGGTCCC
TGCTACCTCGATACAAGGGGTATATCAGATGAGGATGTTCAGAAGCTCAAGGAGGCATACCTCTCAAT
GTCACCCGGTATAATACTCAAGTGGAAGGATGAGAAGATAAACCCCAAGAACACACCCATAGAGATA
TGCGGTTCAGAGCCCTACATAGTTGGTGGTCACGGTCAGGCAGGTTACTGGGTTGATATAAACAGGAA
GACAACACTCGAGGGTCTCTACGCAGCAGGTGATGTTGTTGGTGGTTCACCCAAGAAGTACGTTACAG
GTTGCATGGCAGAGGGTGAGATAGCAGTTGAGGCAGCAATAGAGTACATAAAGTCAATGGAGAACGA
TATAGAGATAGATGAGCAGGAGATAGCAAAGGAGATAGATAGGGTTTTCTACCCCCTCAACAACAAG
AAGGGTGAGTTCTCACCCGATGAGATAGAGGAGAGGATGCAGAAGGTTATGGATGAGTACGCAGGTG
GTATATCATCATACTACAGGGTTAACGAGTCAAAGCTCCTCATAGCAAGGGAGCTCCTCAAGGCAATA
GAGGAGGATCTCTCAAAGATAAAGGTTAGGAACAGGTACGAGCTCATGAAGTACCACGAGGTTGTTG
ATAGGATACTCGTTGCAAGGGCAGTTGTTGAGCACCTCCTCTACAGGAAGGAGACAAGGTGGAAGTG
CTACCAGGAGAGGGTTGATTACCCCGAGATAGATGATAACTGGTTCAAGTTCATAAACTCAAAGTACA
ACTCACAGACAAACGATATAGAGATAATAGAGAGGGAGTACGAGAAGTTCAACCCCGTTATAAACAA
CGATCAC**AGCAGCGGC**CACCACCACCACCACCAC**TGA**GCTAGCATGACTGGTGGACAGCAAATGGG
TCGCGAAGGAGATATACC***ATG***ACAATAAGGATAATAGAGGAGATATGCATAGGTTGCGGTCTCTGCAC
AAAGGTTTGCCCCGGTAACCTCCTCTACCAGAGGGAGGATGGTAAGTCAGAGATAATGGATAAGAGG
GATTGCTGGGATTGCGCAGCATGCGTTAAGGAGTGCCCCGTTAACGCAATAGAGATGTACCTCCAGCC
CGAGATAGGTGGTAGGGGTTCAACACTCAAGGCAAAGAAGACAGATGATTCAATAGTTTGGATAATA
ACAGATAACAACGGTGAGGAGGAGGTTATAGAGGTTAAGAACAAGAAGACATTCGATATG**TGA**GGA
TCC

Red = Insertion of an internal RBS, Blue=linker, **Restriction sites (NcoI and BamHI)**
